# Signaling amplitude molds the *Ras* mutation tropism of urethane

**DOI:** 10.1101/2021.02.09.430515

**Authors:** Siqi Li, Christopher M. Counter

**Affiliations:** Pharmacology and Cancer Biology, Duke University, Durham, North Carolina, USA

## Abstract

RAS genes are commonly mutated in cancers yet despite many possible mutations, cancers have a ‘tropism’ towards a specific subset. As driver mutations, these patterns ostensibly originate from normal cells. High oncogenic RAS activity causes oncogenic stress and different oncogenic mutations can impart different levels of activity. Here we show that changing rare codons to common in the murine *Kras* gene to increase translation shifts tumors induced by the carcinogen urethane from arising from canonical Q_61_ to biochemically less active G_12_ *Kras* driver mutations, despite the carcinogen still being biased towards generating Q_61_ mutations. Loss of p53 to blunt oncogenic stress partially reversed this effect, restoring Q_61_ mutations. Finally, transcriptional analysis revealed similar signaling amongst tumors driven by different mutations and *Kras* alleles. These finding suggest that the RAS mutation tropism of urethane is largely product of selection in normal cells for mutations promoting proliferation without causing oncogenic stress.

**Impact statement:** The bias towards specific *Kras* driver mutations during urethane carcinogenesis appears to arise predominantly from the selection of a narrow window of oncogenic signaling in normal cells.

## Introduction

The RAS genes are mutated in a fifth or more of human cancers (Prior et al., 2020), which is well established to be tumorigenic (Pylayeva-Gupta et al., 2011). There are three individual RAS genes in humans, namely *KRAS, NRAS*, and *HRAS*, three primary sites mutated in human cancers, namely G_12_, G_13_, and Q_61_, and six possible amino acid substitutions at each site arising from a point mutation. As such, there are 54 possible oncogenic mutations, not including rare non-canonical mutations (Hobbs et al., 2016). Mapping these mutations to human cancers reveals a distinct pattern or ‘tropism’ in which specific mutations are characteristics of individual cancer types (Li et al., 2018). The same can be said for other oncogenes, for example, the most frequent EGFR point mutation in glioblastoma is G_598_V/A but L_861_Q in lung adenocarcinoma; the most frequent IDH1 mutation in low-grade glioma is R_132_H but R_132_C in melanoma, and so on (Chang et al., 2016). While these mutational biases are well described, the mechanism responsible is not.

One hint to the mechanism underlying RAS mutation tropism is that oncogenic RAS mutations are thought to occur early. Focusing on lung cancer, in humans oncogenic *KRAS* mutations have been detected in premalignant lesions (Kanda et al., 2012) as well as in multiple regions within the same tumor (Zhang et al., 2014a), indicative an early origin (Wistuba and Gazdar, 2006). In mice, oncogenic *Kras* mutations are capable of initiating pulmonary tumors in carcinogen (McCreery and Balmain, 2017) and genetically engineered (Kwon and Berns, 2013) lung cancer models. As early driver mutations, it follows that RAS mutation tropism is a reflection of the normal cells in which the mutation first occurred. While by its vary classification oncogenic RAS can induce proliferation (Pylayeva-Gupta et al., 2011), high oncogenic signaling through the MAPK effector pathway of RAS can paradoxically induce a stress response in normal cells mediated by the tumor suppressors p16 and p53, which leads to the growth arrest termed senescence (Munoz-Espin and Serrano, 2014). Indeed, hyperactivation of MAPK signaling via the combination of *Kras^G12V^, BrafD^631A^*, and loss of the remaining wildtype *Braf* allele activated p53 and impedes lung tumorigenesis, an effect rescued by pharmacologic inhibition of the MAPK pathway (Nieto et al., 2017).

Accumulating evidence suggests that different oncogenic mutations may impart biochemical differences to the RAS proteins (Munoz-Maldonado et al., 2019; Smith et al., 2013). Relevant to this study, a G_12_D mutation sterically inhibits the catalytic cleft (Parker et al., 2018) while Q_61_R replaces the catalytic amino acid (Buhrman et al., 2010). A direct comparison of the G_12_D mutation to the Q_61_R mutation in Nras revealed the former has lower GTP loading and tumorigenic potential in the skin (Burd et al., 2014) and hematopoietic system (Kong et al., 2016). Indeed, a panel of G_12/13_ Kras mutants introduced by Cas9-mediated gene editing in the lung revealed widely different tumorigenic potentials between different mutants (Winters et al., 2017). Taken together, we hypothesize that the sensitivity of a normal cell to oncogenic signaling amplitude may dictate the type of mutation selected, or to put it another way, selection underpins RAS mutation tropism.

One challenge to testing this hypothesis is trying to backtrack to catch a single, ostensibly random mutagenic event in one gene from one cell, decades before manifesting as cancer in humans. However, in mice the moment of tumor initiation can be precisely defined as the point of carcinogen exposure, with the added benefit that carcinogens model the spontaneous nature of human cancers. Carcinogens can also model RAS mutation tropism. Case in point, the carcinogen urethane found in the fermented foods and alcoholic products (Gowd et al., 2018) primarily induces pulmonary tumors with a very specific Kras^Q61L^ or Kras^Q61R^ driver mutation, depending on the strain (Dwyer-Nield et al., 2010). These oncogenic mutations are also a strong match to the mutation signature of this carcinogen. Urethane-induced mutations conform to the A→T/G consensus sequence derived from comprehensive whole-exome sequencing urethane-induced tumors (Westcott et al., 2015), and the even more restricted C**<u>*A*</u>**N→C**<u>*T/G*</u>**N sequence determined shortly after urethane exposure by different ultra-sensitive sequencing approaches (Li et al., 2020; Valentine et al., 2020), which at the position C**<u>*A*</u>**_182_A gives rise to a *C**<u>T/G</u>**A* mutation encoding Q_61_L/R oncogenic mutations. We thus capitalized on the extreme mutational tropism of urethane for Kras^Q61L/R^-mutant pulmonary tumors to elucidate the effect of a normal cell response on the selection of initiating mutations.

To examine the effect of signaling amplitude on the selection of initiating oncogenic mutations, we genetically manipulated the level of endogenous Kras (to increase oncogenic stress) or the expression of the aforementioned tumor suppressor p53 (to inhibit the cellular response to oncogenic stress) in mice exposed to urethane. In regards to the first genetic change, we compared the native *Kras^nat^* allele that is naturally enriched in rare codons and correspondingly poorly translated (Lampson et al., 2013) to the *Kras^ex3op^* allele, in which 27 rare codons in exon 3 (which is not the site of oncogenic mutations) were converted to common, leading to roughly twice as much Kras protein in the lungs or derived cells of mice (Pershing et al., 2015). In regards to the second genetic change, we evaluated retaining or conditionally inactivating the *Trp53* gene in the lung, which has been shown to suppress oncogenic stress due to high mutant Kras expression in this tissue in vivo (Feldser et al., 2010; Junttila et al., 2010). Using this approach, we show here that the canonical Q_61_L/R mutations are selected against in the more highly expressed *Kras^ex3op^* allele in urethane-induced tumors, even though the carcinogen favors this mutation. Instead, biochemically less active G_12_ mutations are detected, which upon the loss of p53, partially shifts back to Q_61_L/R mutations. p53 loss also promoted the expansion of the normally rare G_12_ mutants in the native *Kras* allele, which was accompanied by an imbalance between the mutant and wildtype transcripts suggestive of higher expression. Taken together, these data support a very narrow window of signaling conducive to tumor initiation, with p53 loss correcting this by suppressing oncogenic stress in more active mutations (Q_61_L/R in *Kras^ex3op^*) and promoting or permitting allelic imbalance of less active mutations (G_12_ mutations in *Kras^nat^*). In agreement, the various combinations of mutations and *Kras* alleles that were tumorigenic induced similar levels of oncogenic signaling, as measured by the transcriptional responses of downstream RAS target genes. In sum, we provide evidence that by modulating endogenous genes in vivo in a model of extreme RAS mutation tropism, selection for a narrow signaling amplitude appears to be the main driving force in the type of Kras driver mutations capable of initiating tumorigenesis.

## Results

### Higher Kras expression accelerates tumorigenesis in the absence of p53

To determine the effect of genetically manipulating oncogene-induced stress on the RAS mutation tropism of urethane, we crossed *Sftpc^CreER/CreER^;Trp53^flox/flox^* mice into a background with two native *Kras* alleles (*Kras^nat/nat^*) or with one (*Kras^nat/ex3op^*) or two (*Kras^ex3op/ex3op^*) copies of the aforementioned *Kras^ex3op^* allele in which rare codons were altered to common in exon 3 (Pershing et al., 2015). The *Sftpc^CreER/CreER^;Trp53^flox/flox^* genotype was chosen as injection of tamoxifen into such mice leads recombination and inactivation of the endogenous *Trp53^flox^* alleles in the type II alveolar cells in the lung (Xu et al., 2012), which is reported to suppress oncogene-induced senescence/apoptosis induced by oncogenic Kras in this organ (Feldser et al., 2010; Junttila et al., 2010). The *Kras^ex3op^* allele was chosen as a way to increase Kras protein expression of the native gene while leaving the rest of the locus almost entirely intact (Pershing et al., 2015). Cohorts of 21-30 mice from each of these three genotypes were injected with tamoxifen to inactive the *Trp53* gene in the lung, followed by exposure to urethane via a single intraperitoneal injection to induce *Kras* mutations. One year later these mice were humanely euthanized, the number and size of tumors determined at necropsy (***Figure 1A*** and ***Supplementary file 1***), and the tumors removed and recombination of the *Trp53^flox^* alleles confirmed by PCR (***Figure 1-figure supplement 1A*** and ***Supplementary file 1***). As a first step, we simply examined the effect of p53 loss on tumorigenesis when Kras expression was altered. In sharp contrast to the previous findings that the *Kras^ex3op^* allele *<u>reduced</u>* urethane carcinogenesis (Pershing et al., 2015), the loss of p53 instead *<u>increased</u>* tumor burden in mice with at least one *Kras^ex3op^* allele (***Figure 1B***), similar to what was observed in a *Cdkn2a* null background (Pershing et al., 2015). This appeared to be a product of more tumors (***Figure 1C***), with a trend towards larger tumors (***Figure 1D***) that reached statistical significance when the data were censored for large (≥ 100 mm^3^) tumors (***Figure 1E***). There was also a trend when the genotypes were subdivided into one or two *Kras^ex3op^* alleles compared to the *Kras^nat/nat^* background, although no difference was observed between *Kras^nat/ex3op^* versus *Kras^ex3op/ex3op^* genotypes (***Figure 1-figure supplement 1B-E***). This suggests that in the absence of p53, the *Kras^ex3op^* allele promotes both the initiation and progression of urethane-induced lung tumors, consistent with p53 suppressing oncogene toxicity to allow oncogenic mutations in the *Kras^ex3op^* allele to exert a more potent signal to drive tumorigenesis.

**Figure 1.**
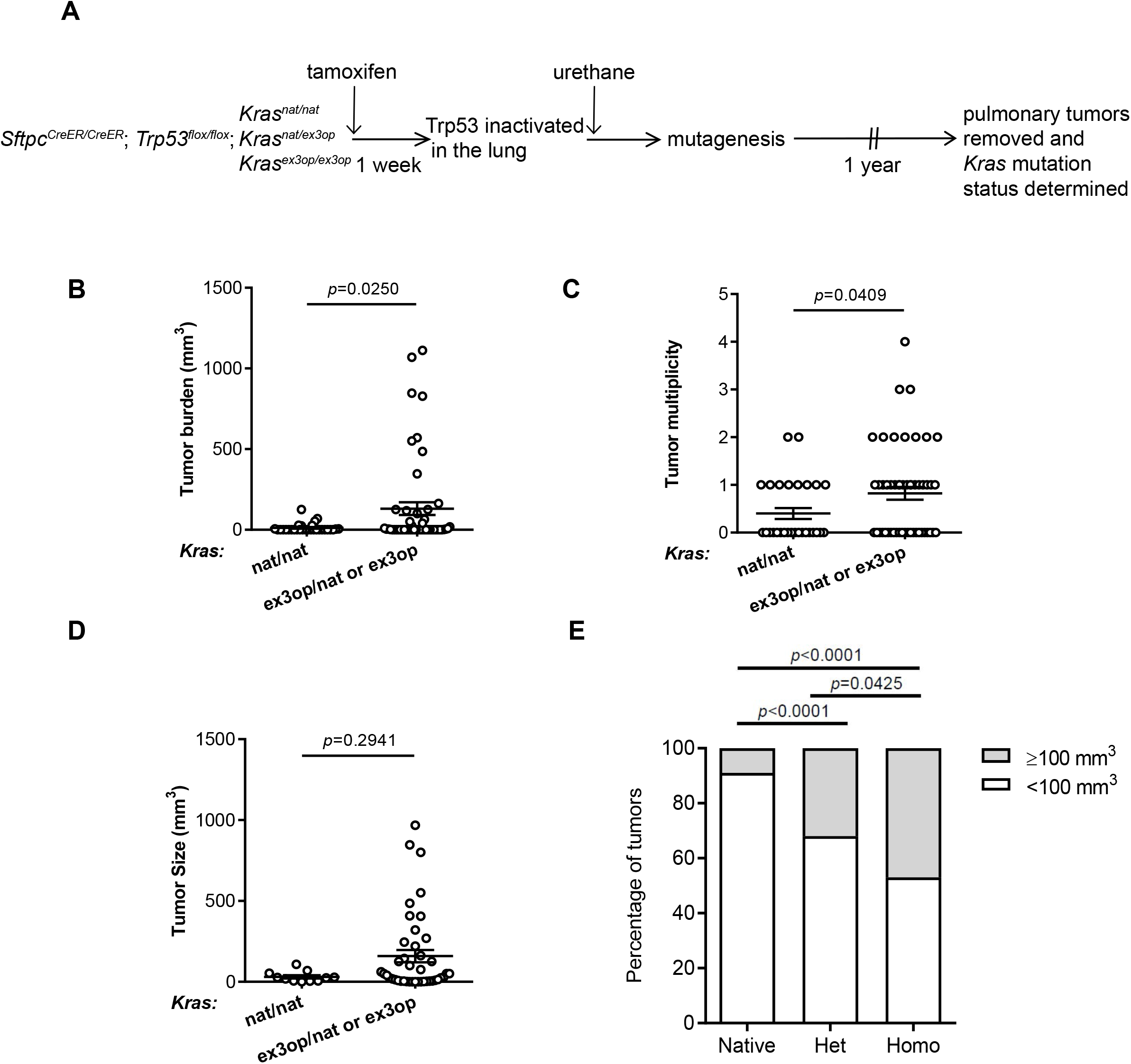
Loss of p53 converts the *Kras^ex3op^* allele from suppressing to enhancing urethane carcinogenesis. (**A**) Experimental design to evaluate the effect of inactivating p53 specifically in the lung on urethane carcinogenesis upon increase Kras expression. (**B-D**) Mean ± SEM of urethane-induced tumor (**B**) burden, (**C**) multiplicity, and (**D**) size in tamoxifen-treated *Sftpc^CreER/ CreER^;Trp53^fl/fl^* mice in a homozygous native ((**B** and **C**) n= 30 mice, (**D**) n=11 tumors) and heterozygous or homozygous ((**B** and **C**) n= 51 mice, (**D**) n=42 tumors) ex3op Kras background. Mann-Whitney test. (**E**) % of tumors ≥ (grey bar) or < (white bar) 100 mm^3^ in tamoxifen-treated *Sftpc^CreER/CreER^;Trp53^fl/fl^* mice in *Kras^nat/nat^* (n=11 tumors), *Kras^ex3op/nat^* (n=25 tumors), or *Kras^ex3op/ex3op^* (n=17 tumors) background after urethane exposure. Twosided Fisher’s exact test.

### Loss of p53 reprograms the extreme RAS mutation tropism of urethane

To specifically address the effect of genetically inactivating the *Trp53* gene on the type of oncogenic mutations arising in the native *Kras^nat^* versus the codon-optimized *Kras^ex3op^* allele in tumors induced by urethane, we compared the *Kras* mutation status in tumors from the above *Sftpc^CreER/CreER^; Trp53^flox/flox^; Kras^nat/ex3op^* mice injected with tamoxifen (termed *Trp53^-/-^*), as the heterozygous status of the *Kras^nat/ex3op^* background allows for the most direct comparison, to tumors from a parallel control cohort not injected with tamoxifen (termed *Trp53^+/+^*) prior to urethane exposure (***Figure 2A*** and ***Supplementary file 1***). As mentioned above, the *trp53^flox^* alleles the were confirmed by PCR to be recombined in tumors from the *Trp53^-/-^* background. The same analysis was performed on tumors from the *Trp53^+/+^* cohort, which identified one tumor having a significant degree *Trp53^flox^* recombination, which was excluded from the analysis of comparing *Trp53^+/+^* versus *Trp53^-/-^* mice (***Figure 2-figure supplement 1A***). Consistent with the role of p53 as a tumor suppressor during lung tumor progression (Feldser et al., 2010; Junttila et al., 2010), loss of p53 tracked with larger, but not more tumors (***Figure 2B*** and ***Figure 2-figure supplement 1B,C***). To examine the effect of *p53* deficiency on the RAS mutation tropism of urethane, we sequenced *Kras* derived from mRNA isolated from these lung tumors and screened for mutations at the three main hotspots of G_12_, G_13_, and Q_61_. In complete agreement with the previous observation that the increased protein expression of the endogenous *Kras^ex3op^* allele shifts the RAS mutation tropism of urethane from the canonical Q_61_ to G_12_ oncogenic mutations (Pershing et al., 2015), tumors with an oncogenic mutation in the *Kras^ex3op^* allele from control *Trp53^+/+^* mice similarly had G_12_ oncogenic mutations in this allele (***Figure 2C***). Having established that urethane behaves identically as previously reported in this regard, we turned our attention to the types of mutations recovered in the *Kras^ex3op^* allele from the urethane-induced tumors of the *Trp53^-/-^* mice. Sequencing revealed that 40% hotspot oncogenic mutations in the *Kras^ex3op^* allele were now detected at Q_61_ (***Figure 2C***), or to put it another way, the loss of p53 shifted the mutations in the *Kras^ex3op^* allele back to more of the canonical Q_61_ mutations of urethane.

**Figure 2.**
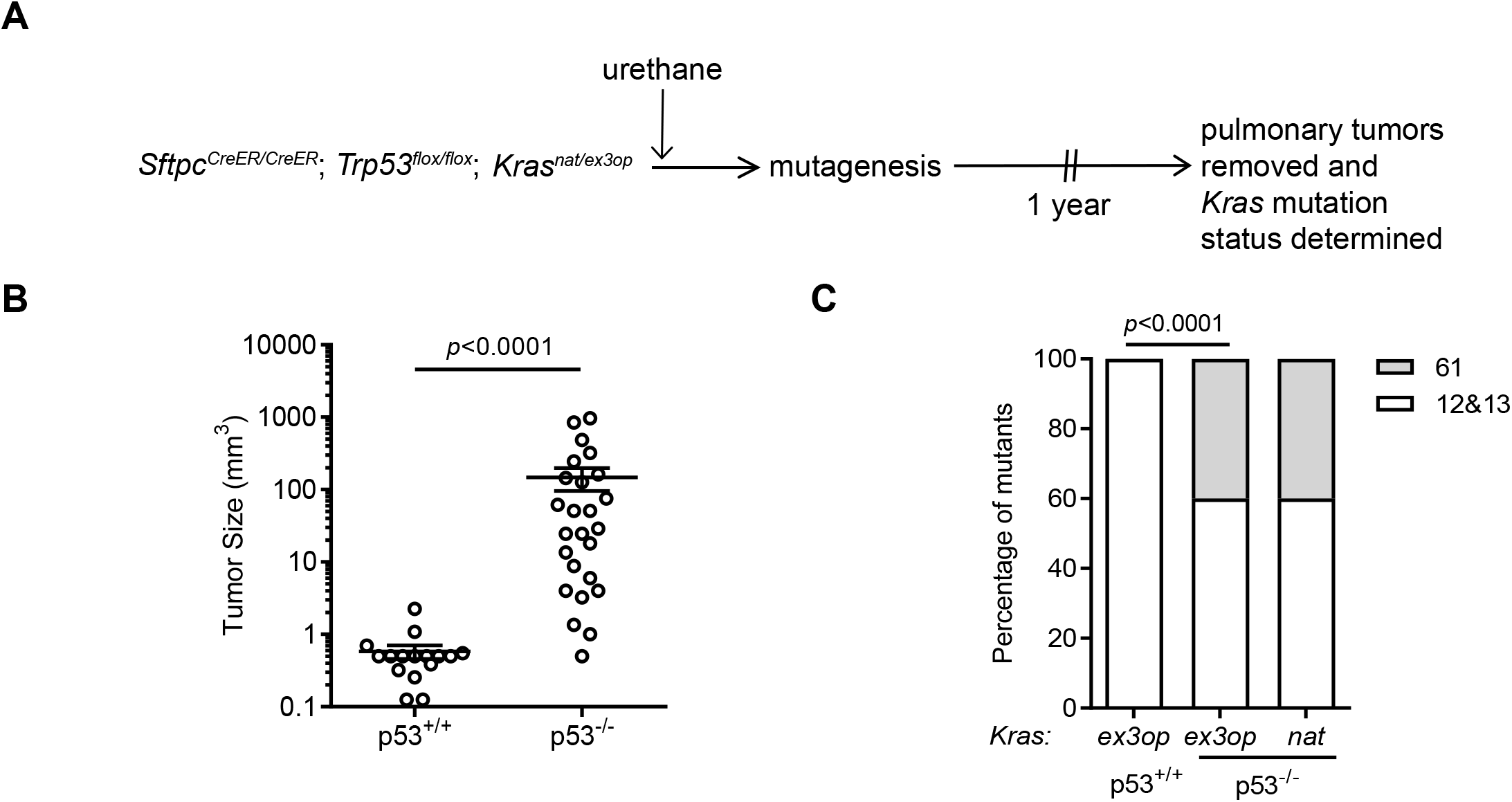
Loss of p53 reprograms the RAS mutation tropism of urethane. **(A)** Experimental design to obtain urethane-induced lung tumors from p53^+/+^ mice. **(B)** ± SEM of urethane-induced tumor size in *Sftpc^CreER/CreER^;Trp53^fl/fl^;Kras^ex3op/nat^* mice not treated (p53^**+/+**^, n=16 tumors) or treated with tamoxifen (p53^**-/-**^, n=25 tumors). Mann-Whitney test. **(C)** % of urethane-induced tumors with an oncogenic mutation at codon G12/13 (white bar) versus Q61 (grey bar) in the *Kras^nat^* versus *Kras^ex3op^* allele in *SftpC^CreER/CreER^;Trp53^fl/fl^;Kras^ex3op/nat^* mice not treated (p53^+/+^) or treated with tamoxifen (p53^**-/-**^) where indicated. n=4 tumors ex3op p53^+/+^, 5 tumors nat p53^-/-^ and 10 tumors ex3op p53^-/-^. Two-sided Fisher’s exact test.

### The mutation signature of urethane is not affected by the *Kras^ex3op^* allele

To determine if the shift in oncogenic mutations from Q_61_ in the *Kras^nat^* allele to G_12_ in the *Kras^ex3op^*, and then back again in the *Trp53^-/-^* background, resides at the level of the locus or with the amount of encoded protein, we tested whether urethane induces different mutations in the *Kras^ex3op^* versus *Kras^nat^* allele. We thus turned to the ultra-sensitive Maximum Depth Sequencing (MDS) assay (Jee et al., 2016), which we adapted for the mammalian genome to detect urethane-induced mutations within days of carcinogen exposure (Li et al., 2020). Since only a short region of genomic DNA can be sequenced by this approach, it is not possible to track oncogenic mutations in exon 1 or 2 and also determine the identity of the *Kras* allele (*native* versus *ex3op*) based on the codon usage in exon 3. Thus, we compared mutations arising in the *Kras^nat/nat^* versus *Kras^ex3op/ex3op^* genotype. To ensure potent mutagenesis for detection purposes, these two strains were crossed into the pure 129 background, which is sensitive to urethane (Malkinson and Beer, 1983; Shimkin and Stoner, 1975), and injected three times instead of just once as above with either the vehicle PBS or urethane. Seven days later, before overt cell selection (Li et al., 2020), the mice were humanely euthanized and their lungs removed and subjected to MDS sequencing to determine both the mutation signature and the type of *Kra*s driver mutations induced by urethane (***Figure 3A*** and ***Supplementary file 2***). Mutation frequencies based on MDS sequencing of *Kras* exon 1 and exon 2 were averaged for A→T transversions, log_10_ transformed, and displayed in a heatmap format. This revealed a trend towards A→T transversions within the context of a 5’ C in both the native and ex3op alleles of *Kras* specifically in urethane-exposed mice (***Figure 3B***), consistent with previously identified bias for urethane (Li et al., 2020; Valentine et al., 2020; Westcott et al., 2015). Moreover, the frequency of these *C**<u>A</u>**→C**_T_*** mutations, which give rise to Q_61_L (CA_182_A→C**<u>*T*</u>**A), are significantly higher than *G**<u>G</u>**→G**<u>A</u>*** mutations, which give rise to G_12_D (GG35T→G**<u>*A*</u>**T) and G_13_D (GG38C→G**<u>*A*</u>**C), in both *Kras^nat/nat^* and *Kras^ex3op/ex3op^* mice (***Figure 3C***). Similar results were found upon repeating the experiment with a single injection of urethane in the less sensitive 129/B6 mixed strain background using *Sftpc^CreER/CreER^;Trp53^flox/flox^;Kras^nat/nat^* versus *Sftpc^CreER/CreER^; Trp53^flox/flox^;Kras^ex3op/ex3op^* mice treated or not with tamoxifen (***Figure 3-figure supplement 1A***). The only exception was that far fewer *C**A**→C**Ţ*** mutations were detected in general, and perhaps as a consequence, the difference between the frequency of *C**<u>A</u>**→C**<u>T</u>***and *G**<u>G</u>**→G**<u>A</u>*** mutations was no longer significant, although *G**<u>G</u>**→G**<u>A</u>*** mutations were detected in both the urethane and PBS cohorts, suggesting false positives, although other interpretations are possible (***Figure 3-figure supplement 1B,C***). Thus, with the above proviso, urethane mutagenesis does not appear to be changed at the *Kras^ex3op^* locus, at least within the detection limit of the MDS assay. These and the above findings support that the observed bias towards G_12/13_-driver mutation in the *Kras^ex3op^* allele in urethane-induced tumors is a product of negative selection against the biochemically more active Q_61_ oncogenic mutation, rather than a change in the mutational spectrum of urethane. In other words, oncogenic stress molds the type of RAS mutations conducive to initiate tumorigenesis irrespective of the mutational signature.

**Figure 3.**
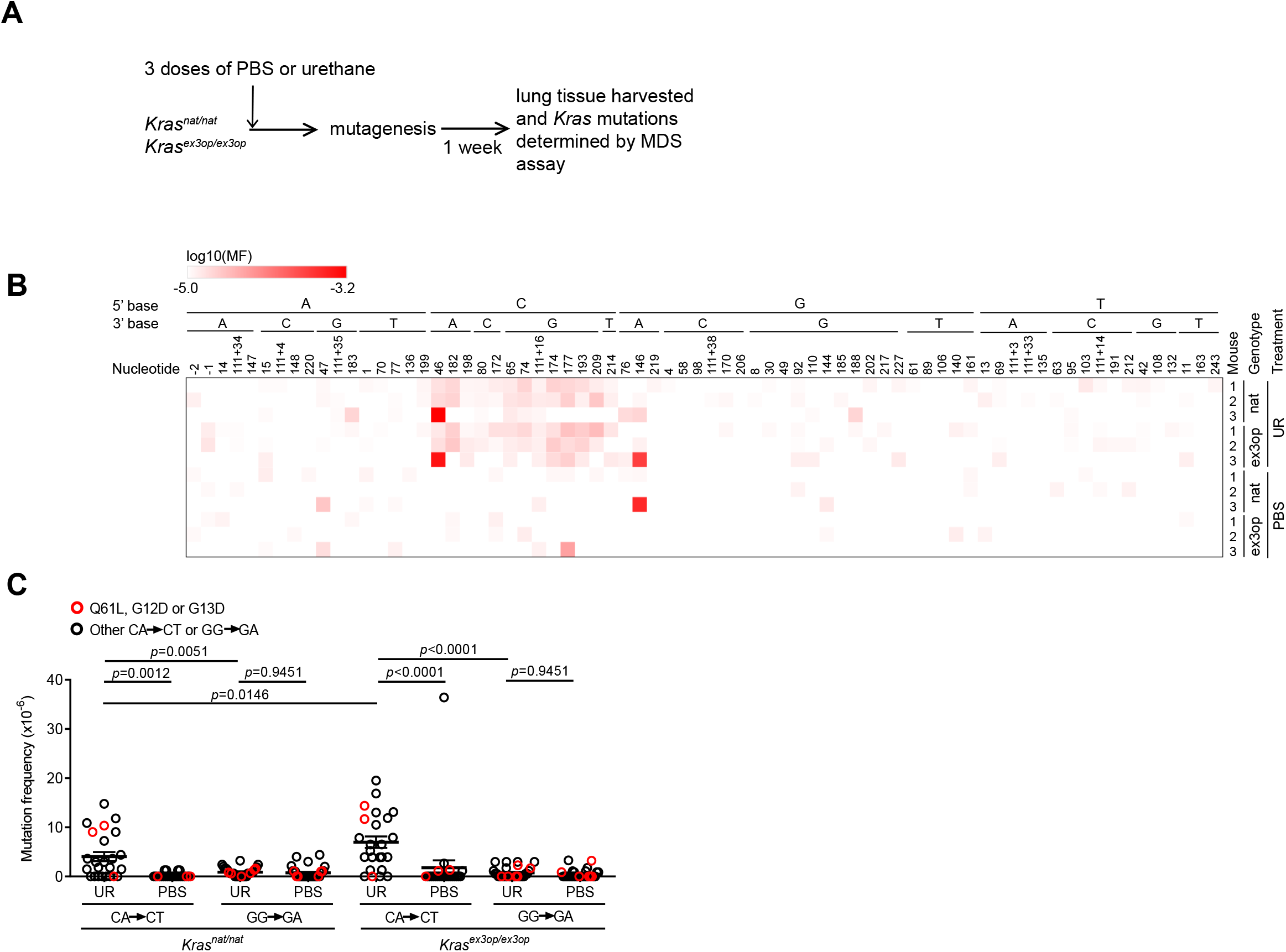
The mutation signature of urethane is not affected by the *Kras^ex3op^* allele. **(A)** Experimental design to identify mutations induced by urethane in mouse lung in a *Kras^nat^* versus *Kras^ex3op^* background. **(B)** Heatmap of the log transformed mutation frequency (MF) of A->T transversions determined by MDS sequencing the exon 1 and exon 2 of *Kras* in from the lungs of mice exposed to urethane (UR) in a *Kras^nat/nat^* (nat) (n=3 mice) versus *Kras^ex3op/ex3op^* (ex3op) (n=3 mice) background. Nucleotide number as well as the 5’ and 3’ base of the substituted A are shown at the top; “-” indicates nucleotides upstream of ATG start codon in 5’UTR; “111+” indicates nucleotides in the intron downstream of exon 1. **(C)** Mean ± SEM mutation frequency of all CA-> CT mutations in *Kras* exon 2, with Q61L mutation highlighted in red, as well as all GG-> GA mutations in *Kras* exon 1, with G12D and G13D muations highlighted in red, derived from the aforementioned MDS sequencing of *Kras* exon 1 and 2 from the lungs of *Kras^nat/nat^* versus *Kras^ex3op/ex3op^* mice treated with either urethane or PBS (n=3 mice each). Holm-Sidak multiple comparisons test following one-way ANOVA.

### p53 loss tracks with an mRNA allelic imbalance of less active oncogenic mutations

Not only was there was a shift in the oncogenic mutations in the *Kras^ex3op^* allele upon the loss of p53, as noted above, but surprisingly also in the *Kras^nat^* allele. In more detail, we found that 60% hotspot mutations in the *Kras^nat^* allele of urethane-induced tumors from the *Sftpc^CreER/CreER^;Trp53^flox/flox^; Kras^nat/ex3op^* mice in which the *Trp53* gene was recombined in the lung occurred at codon G_12_, and in one case also G_13_ (***Figure 2C***). We also note that the percentage of tumors that have *Kras* hotspot mutations is higher in p53^-/-^ mice (***Figure 4-figure supplement 1A***). In addition, even though oncogenic mutations are generally more frequent in the *Kras^ex3op^* allele, the percentage of tumors with mutations in the native allele is higher in p53^-/-^ mice (***Figure 4-figure supplement 1B***). One interpretation of these findings is that the absence of p53 enhances the ability of G_12/13_ mutations to be productive. Given the above tight relationship between Kras expression and mutation type, we explored a possible relationship between p53 loss and higher expression of *Kras* alleles with a G_12/13_ mutation. To this end, we calculated the ratio of mutant to wildtype *Kras* mRNA based on the number of cDNA sequencing reads matched to the mutant or wildtype allele from the above analysis. This revealed a clear demarcation in *Kras* mRNA levels between those alleles with a Q_61_ versus a G_12/13_ oncogenic mutation. In detail, a waterfall plot revealed the mutant:wildtype ratio was higher in G_12/13_-mutant *Kras* alleles, with an average ratio of ~2 copies for the mutant *Kras* transcript to each copy of wildtype *Kras* counterpart. Conversely, the mutant:wildtype ratio was lower in Q_61_-mutant *Kras* alleles, with an average ratio of ~0.8 copies for the mutant *Kras* transcript to each copy of wildtype *Kras* counterpart (***Figure 4A*** and ***Figure 4-figure supplement 1C,D***). The allelic imbalance in the G_12/13_-mutant tumors appears to be important for tumorigenesis. Namely, cross referencing the mutant:wildtype *Kras* ratio to the mutation type and size of tumors revealed that G_12/13_-mutant tumors with a mutant:wildtype ratio ≥ 1.5 are larger, reaching the same size of Q_61_-mutant tumors (***Figure 4B***). This suggests that to achieve optimal signaling amplitude, *Kras* with less active G_12/13_ oncogenic mutations undergo a selection for higher expression to compensate for the lower signaling, while if anything, the reverse was seen for more active Q_61_ oncogenic mutations.

**Figure 4.**
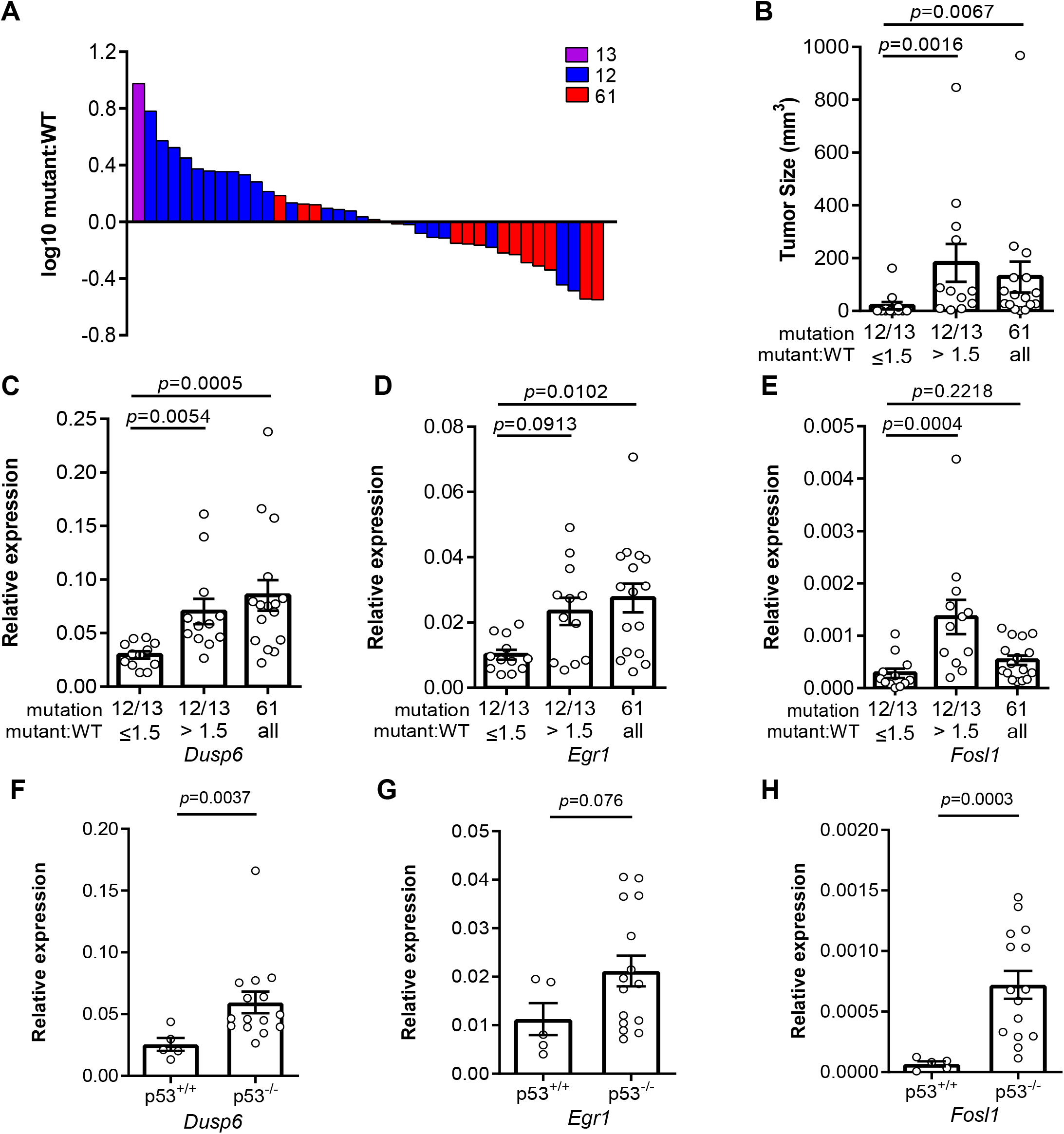
Loss of p53 promotes higher expression of weaker oncogenic mutations. **(A)** Log1O-transformed ratio of mutant to wildtype *Kras* mRNA determined by RT-qPCR in all Kras hotspot-mutant tumors (n=40) derived from figures 1 and 2. **(B)** Mean ± SEM size of tumors with a G_12/13_ oncogenic Kras mutation with a high (>1.5, n=12 tumors) versus low (≤ 1.5, n=12 tumors) mutant:WT ratio versus tumors with a Q61 oncogenic Kras mutation (n= 16 tumors). Dunn’s multiple comparison test following Kruskal-Wallis test. (**C-H**) Mean ± SEM levels of the indicated mRNAs normalized to β-actin (Relative expression) in (**C-E**) tumors with a G12/13 oncogenic Kras mutation with a high (>1.5, n=12 tumors) versus low (≤ 1.5, n=12 tumors) mutant:WT ratio versus tumors with a Q61 oncogenic Kras mutation (n=16 tumors) or (**F-H**) tumors from *Sftpc^CreER/CreER^;Trp53^fl/fl^;Kras^ex3op/nat^* mice not treated (p53^+/+^, n=5 tumors) or treated with tamoxifen (p53^**-/-**^, n=15 tumors) partitioned by p53 mutation status. (**C-E**) Dunn’s multiple comparison test following Kruskal-Wallis test. (**F-H**) Mann-Whitney test.

### Allelic mRNA imbalance of G_12/13_-mutant *Kras* alleles tracks with Ras signaling

To test the possibility that there is a selection for a specific signaling amplitude, we measured the degree of oncogenic RAS signaling through the MAPK pathway, the very pathway promoting proliferation (Drosten and Barbacid, 2020; Hymowitz and Malek, 2018; Ryan et al., 2015) or senescence (Munoz-Espin and Serrano, 2014), by assaying the transcriptional level by qRT-PCR of the three well-described downstream target genes *Dusp6* (Buffet et al., 2017; Kidger and Keyse, 2016; Unni et al., 2018; Zhang et al., 2010), *Egr1* (Esnault et al., 2017; McMahon and Monroe, 1995; Swarbrick et al., 2008), and *Fosl1* (Chung et al., 2017; Esnault et al., 2017; Gillies et al., 2017; Vallejo et al., 2017a; Vallejo et al., 2017b). Binning the relative expression of these three genes into *Kras* G_12/13_-mutant tumors with a high (> 1.5) versus low (≤ 1.5) mutant:wildtype ratio revealed that *Kras* G_12/13_-mutant tumors with a high ratio expressed of these three genes more strongly than those with a low ratio. Furthermore, when we compared the relative expression of these three genes in *Kras* Q_61_-mutant tumors, *Dusp6* and *Egr1* were expressed higher compared to the *Kras* G_12/13_-mutant tumors with a low ratio (***Figure 4C-E***). Plotting the signaling intensity against the mutant:wildtype cDNA ratio of individual tumors showed that signaling intensity is general correlated with the allelic ratio for G_12/13_ mutations (***Figure 4-figure supplement 1E-G***). As such, the increase in expression of the G_12/13_-mutant *Kras* alleles, as measured by a high mutant:wildtype ratio, manifests as an increase in Ras signaling, matching that achieved by the more potent Q_61_-mutated *Kras* alleles. To assess the effect of p53 on this relationship, we compared *Dusp6*, *Egr1*, and *Fosl1* expression between tumors with and without p53. This revealed higher expression of all three genes in the latter tumors, consistent with the loss of p53 permitting higher Ras signaling (***Figure 4F-H***). p53 loss also tracked with higher mutant:wildtype allelic ratio in tumors with *Kras* G_12/13_-mutations in the *Kras^ex3op^* allele (***Figure 4-figure supplement 1H***). Finally, we compared *Dusp6, Egr1*, and *Fosl1* expression in tumors of different sizes, which revealed higher Ras signaling in larger tumors, regardless of how this achieved (***Figure 4-figure supplement 1I-K***). Collectively, these findings suggest that the absence of p53 expands the spectrum of driver *Kras* mutations to include less active mutations by permitting or even fostering an increase in their expression; once again favoring a model whereby an optimal oncogenic amplitude is selected to initiate tumorigenesis.

## Discussion

Here we show that p53 loss reprograms RAS mutation tropism. On one hand, loss of this tumor suppressor shifts the canonical Q_61_ oncogenic mutations normally detected the native *Kras^nat^* allele of urethane-induced tumors towards the usually rare G_12/13_ mutations in conjunction with an increase in the mutant:wildtype ratio. On the other hand, the loss of p53 shifts the prevalence of G_12/13_ mutations in the codon optimized and more highly translated *Kras^ex3op^* allele towards Q_61_ mutations. The one common theme in both these shifts is oncogenic amplitude, which argues that the degree of oncogenic signaling dictates the type of mutation conducive to initiate tumorigenesis (***Figure 5***). In agreement, oncogenic signaling, as measured by the expression of known RAS downstream target genes, was similar between *Kras^ex3op^* with Q_61_ mutations and *Kras^nat^* with G_12/13_ mutations exhibiting an increase in the mutant:wildtype ratio, suggesting that expression difference render Q_61_ and G_12/13_ mutations fungible. While there is no question that the level of oncogenic signaling can influence tumorigenesis, what is new here is that the sensitivity of normal cells to oncogenic signaling underlies the *<u>type</u>* of oncogenic mutations selected, which is relevant to the RAS mutation tropism of human cancers. We also note that neither the loss of p53 nor the change in codon usage in *Kras* altered the mutation signature of urethane, arguing that a selection for an optimal oncogenic signal supersedes the mutational bias of the urethane. Such a finding speaks to the somewhat complexing discordance often observed between the mutation signature of tumors and the corresponding driver mutation in human cancers (Dietlein et al., 2020; Temko et al., 2018), even in cancers in which the mutational signature can be ascribed to a specific mutagenic process (Buisson et al., 2019).

**Figure 5.**
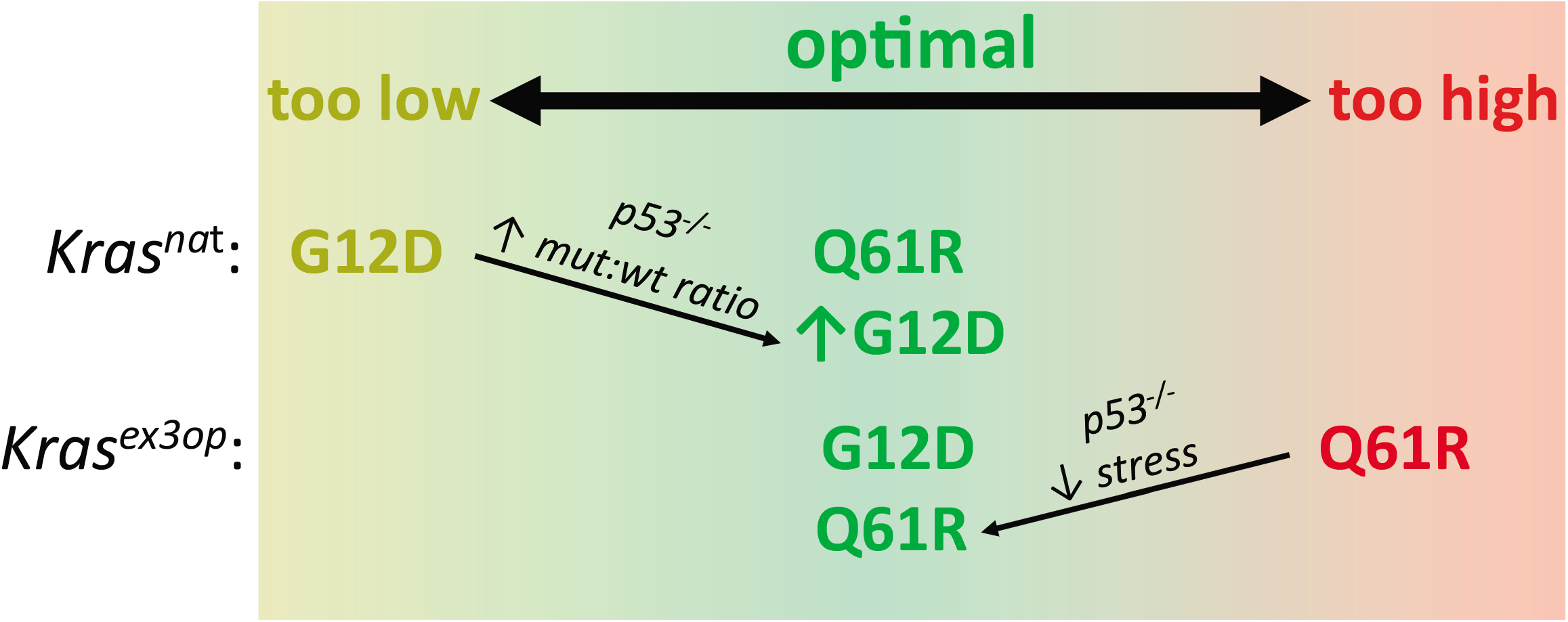
Optimal signaling is required for effective tumor initiation. Signaling from a G12D mutation in the native (nat) *Kras* allele and from a Q61R mutation in the codon optimized (ex3op) *Kras* allele are outside of the window of optimal signaling amplitude achieve by *Kras^nat^*(G12D) and *Kras^ex3op^*(Q61R). Loss of p53 alleviates the selection against oncogenic stress and allows the recovery of a Q61R mutation in *Kras^ex3op^* allele or a G12D mutation in the *Kras^nat^* allele with elevated mutant:wildtype (mut:wt) mRNA allelic ratio.

There are four suppositions to the above model (***Figure 5***). First, Q_61_ oncogenic mutations in Kras activate the MAPK pathway more potently than G_12/13_ mutations. In agreement, a Q_61_R mutation in Nras and/or Kras have a much (1,000 times) slower intrinsic hydrolysis rate, higher GTP-loading, and more robust activation of the MAPK pathway, and when activated in the skin or hematopoietic system in vivo, are more tumorigenic than the corresponding G_12_D mutation (Burd et al., 2014; Kong et al., 2016; Pershing et al., 2015). We also find that the expression of downstream Ras target genes is higher in tumors with a Q_61_ mutation compared to G_12/13_ mutation in the absence of an allelic imbalance. Second, increasing Kras level increases signaling. In agreement, the *Kras^ex3op^* allele has been documented to express more protein in the lung (Pershing et al., 2015) and increase ERK phosphorylation in hematopoietic stem cells (Sasine et al., 2018). Altering rare codons to common in ectopic or endogenous human *KRAS* also leads to higher translation, KRAS protein expression, and/or MAPK activation (Ali et al., 2017; Fu et al., 2018; Lampson et al., 2013; Pershing et al., 2015; Peterson et al., 2020). Nevertheless, it remains possible that rare codons may be differentially translated (Quax et al., 2015) or higher Kras protein expression reduces MAPK signaling through negative feedback (Shin et al., 2009) specifically in the cell-of-origin of these pulmonary tumors. Third, high oncogenic signaling is a negative selection, and fourth, this selection is p53-dependent; two points well supported in multiple model systems (Munoz-Espin and Serrano, 2014). As such, it is reasonable to propose that oncogenic amplitude is a driving force in the selection of type of oncogenic Kras mutations initiating tumorigenesis (***Figure 5***).

We note that urethane has a bias towards C**<u>*A*</u>**N→C**<u>*T/G*</u>**N mutations that is absent in both strand orientations at codons encoding G_12_ and G_13_, and the mutagenic signature of urethane was the same in the *Kras^nat^* and *Kras^ex3op^* alleles. This suggests that G_12/13_ oncogenic mutations are extremely rare, with urethane either inducing these mutations directly at a low frequency, or inducing a cooperating mutation or non-mutational event that stimulates a rare, pre-existing Kras-mutant cell to expand. In either scenario, the very fact that tumors arise with G_12/13_ oncogenic mutations argues for potent selection of an extremely rare oncogenic mutation over the far more common mutations favored by the carcinogen. Furthermore, if these mutations are indeed pre-existing, such a finding elevates the importance of promotional events (secondary mutations, inflammation, etc.) in the process of tumor initiation (Balmain, 2020). We also note an imbalance in the mRNA ratio of the mutant:wildtype *Kras* alleles in some urethane-induced tumors. Amplification of oncogenic *Kras* alleles has been previously documented in various mouse cancer backgrounds (Chung et al., 2017; Mueller et al., 2018; Westcott et al., 2015). What is new, however, is that this tracked with the *<u>type</u>* of oncogenic mutation, and even more interesting, in opposite directions. Namely we observed a bias towards a high mutant:wildtype mRNA ratio in G_12/13_-mutant *Kras* alleles in the absence of p53. Few tumors show amplification of the G_12/13_ mutant allele based on genomic copy number analysis (***Figure 4-figure supplement 2***), pointing towards transcriptional or post-transcriptional mechanisms. One interpretation of these findings is that loss of p53 allows expansion of tumors with G_12/13_-mutant *Kras* alleles that are more highly expressed, an observation borne out by signaling analysis. Increasing MAPK signaling by paradoxical activation with a BRAF inhibitor expands the number of cell types that are tumorigenic upon activation of the *LSL-Kras^G12D^* allele in the lung (Cicchini et al., 2017), hence it is quite possible that a similar phenomenon is at play here as well. Such higher expression could be a product of natural variation of gene expression, a mutation caused by urethane, or p53 loss itself. We also observed a bias towards a low mutant:wildtype mRNA ratio in Q_61_-mutant *Kras* alleles. Such a finding is consistent with this mutation inducing a strong oncogenic signal, and hence a selection against increased expression. Interestingly, this ratio was low irrespective of the p53 status. Perhaps there is an upper limit to oncogenic signaling, even in the absence of p53, that is tumor-promoting. In support, we note that G_12/13_-mutant tumors with a high mutant:wildtype mRNA ratio had the same Ras signaling output as Q_61_-mutant tumors, at least as assessed by the expression level of genes downstream of Ras. Indeed, RAS-mutant human tumors, which often have disruption of p53 and other tumor suppressors, typically do not have BRAF or EGFR mutations and vice versa outside of drug resistance settings (Sanchez-Vega et al., 2018). Moreover, co-expressing oncogenic RAS with oncogenic BRAF or EGFR can be growth suppressive (Cisowski et al., 2016; Unni et al., 2018).

In conclusion, multiple independent lines of investigation support a model whereby the sensitivity of a normal cell to an narrow range of oncogenic signaling dictates the nature of mutation conducive to tumor initiation. Too low, a mutation fails to induce proliferation, as supported by the finding that less active G_12_ mutants are rarely recovered in the native *Kras* allele unless accompanied by an allelic imbalance permitted or fostered by the absence of p53. Too high, a mutation induces arrest, as supported by the finding that more active Q_61_ mutants are not recovered from the more highly expressed *Kras^ex3op^* allele unless oncogene-induced senescence is suppressed by the loss of p53 (***Figure 5***). Moreover, this selection pressure far exceeds the mutational bias of the mutagen, as supported by the observations that G_12_ mutations fail to match the urethane mutational signature, yet nevertheless arise at a high frequency in urethane-induced tumors in a *Kras^ex3op^* background. Such findings speak directly to the extreme mutational bias of urethane-induced tumors, and perhaps more broadly, provides mechanistic insight into RAS mutation tropism in human cancers.

## Materials and Methods

### Key Resource Table

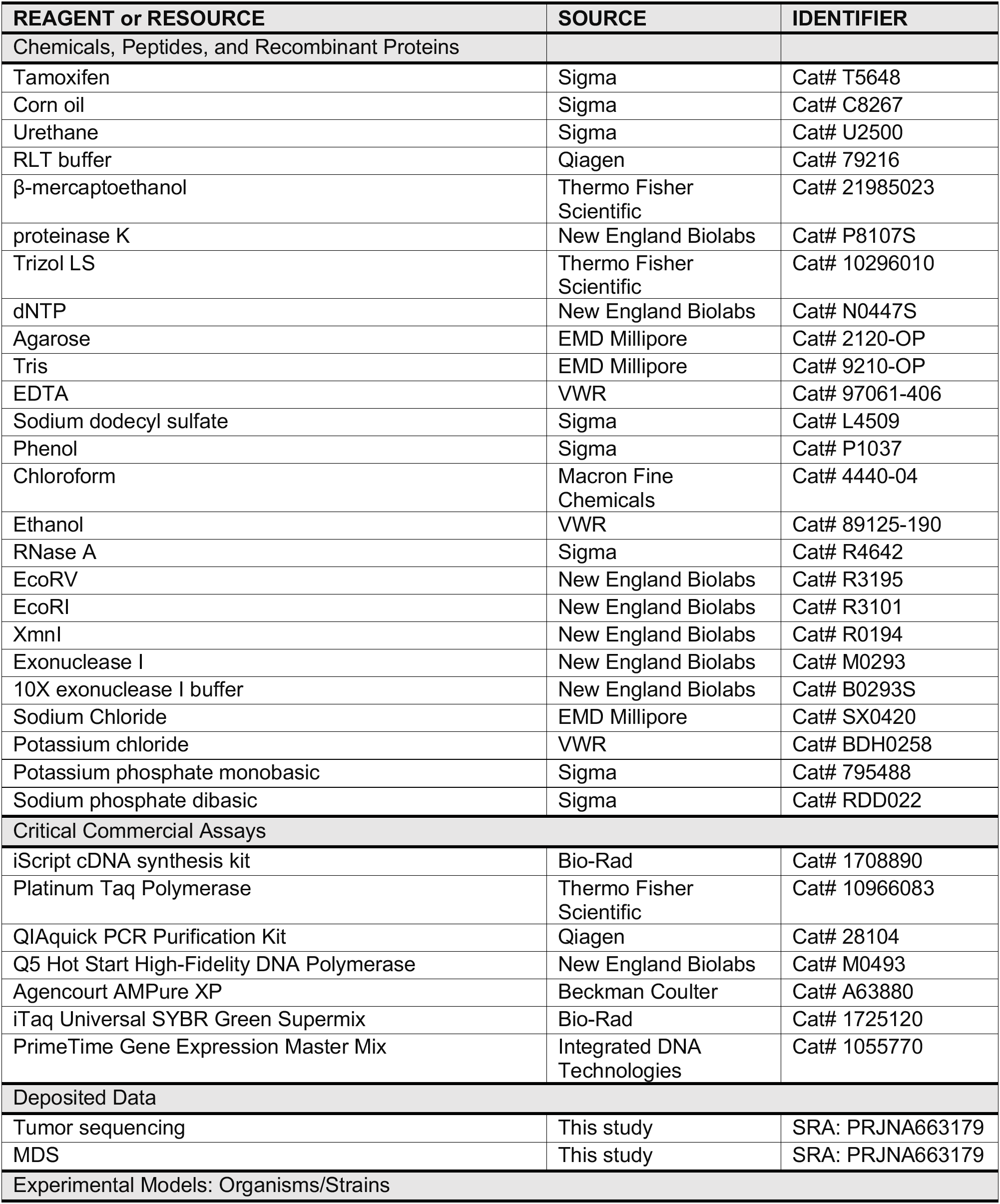
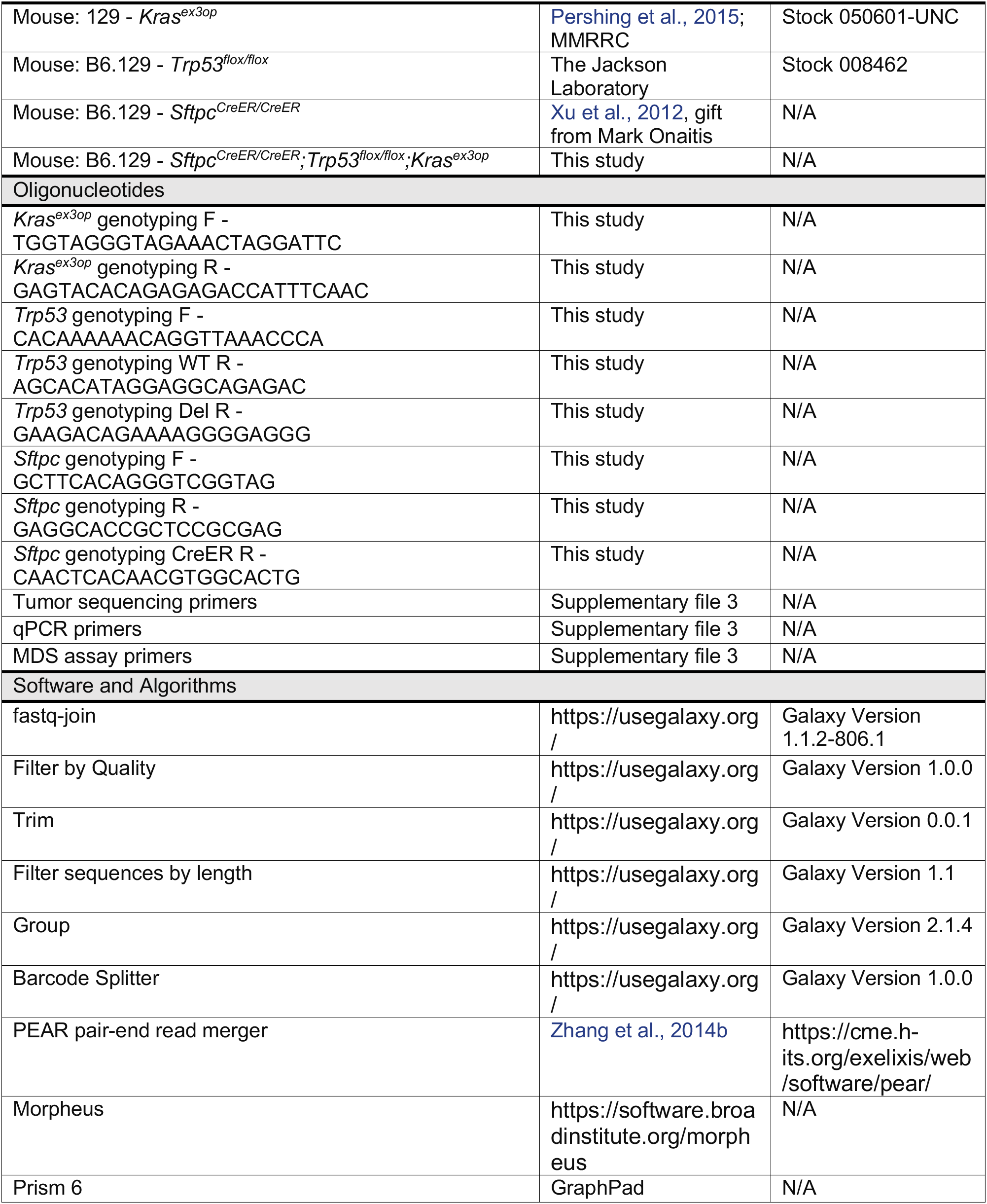

### Mice

*Kras^ex3op^* mice were previously described (Pershing et al., 2015), *Trp53^flox/flox^* mice (Marino et al., 2000) were purchased from The Jackson Laboratory (Jax # 008462), and *Sftpc^CreER/CreER^* mice (Xu et al., 2012) were kindly provided by Mark Onaitis (University of California at San Diego). *Sftpc^CreER/CreER^;Trp53^flox/flox^; Kras^ex3op^* mice were from a mixed 129 x C57BL/6 background. *Kras^ex3op^* mice used for mutagenesis studies were from a pure 129 background. Mice were genotyped using the following primers:

*Kras native* and *ex3op* alleles:

*Kras* F: 5’-TGGTAGGGTAGAAACTAGGATTC-3’
*Kras* R: 5’-GAGTACACAGAGAGACCATTTCAAC-3’
Products: 504 bp (*Kras^ex3op^*) or 614 bp (*Kras^nat^*)

*Sftpc^CreER^* allele:

*Sftpc* F: 5’-GCTTCACAGGGTCGGTAG-3’
*Sftpc* R: 5’-GAGGCACCGCTCCGCGAG-3’
*Spc-CreER* R: 5’-CAACTCACAACGTGGCACTG-3’
Products: 550 bp (*Sftpc*) or 500 bo (*Sftpc^CreER^*)

*Trp53* allele:

*Trp53* F: 5’-CACAAAAAACAGGTTAAACCCA-3’
*Trp53* WT R: 5’-AGCACATAGGAGGCAGAGAC-3’
*Trp53* Del R: 5’-GAAGACAGAAAAGGGGAGGG-3’
Products: 288 bp (wildtype *Trp53*), 370 bp (unrecombined *Trp53fl*), and 612 bp (recombined *Trp53fl*).

All animal experiments were approved by Duke IACUC.

### Tumor experiments

A 20 mg/ml tamoxifen (Sigma T5648) solution was made by dissolving tamoxifen in corn oil. Six-to eight-week-old mice of the indicated genotypes were injected intraperitoneally with 0.25 mg/g body weight tamoxifen every other day for four doses. Urethane (Sigma U2500) was dissolved in PBS and injected intraperitoneally at dose of 1 mg/g body weight 1 week after the last injection of tamoxifen. ~12 months after urethane injection, all mice were humanely euthanized, after which lung tumors were counted, measured, and microdissected for RNA and DNA extraction. The 12 month time point was chosen based on similar previous studies (Dwyer-Nield et al., 2010; Gurley et al., 2015; Miller et al., 2003; You et al., 1989) but also to take into account a potentially longer tumor latency in the mixed 129 x C57BL/6 background (Malkinson and Beer, 1983; Shimkin and Stoner, 1975). Tumor volume was calculated as ½ (length × width^2^). Tumor burden was calculated as the sum of tumor volumes.

### Mutagenesis experiments

Six-to eight-week-old *Kras^nat/nat^* or *Kras^ex3op/ex3op^* mice were intraperitoneally injected daily for three days with either urethane or PBS as above. These mice were humanely euthanized 1 week later and the lungs collected for the extraction of genomic DNA. Alternatively, six-to eight-week-old *Sftpc^CreER/CreER^;Trp53^flox/flox^;Kras^nat/nat^* or *Sftpc^CreER/CreER^;Trp53^flox/flox^;Kras^ex3op/ex3op^* mice were or were not injected intraperitoneally with 0.25 mg/g body weight tamoxifen every other day for four doses. One week after the last injection of tamoxifen, the same mice were intraperitoneally injected with one dose of either urethane or PBS. The mice were humanely euthanized 1 month later and the lungs collected for the extraction of genomic DNA.

### RNA and DNA extraction

Tumors or normal lung tissues were lysed in RLT buffer (Qiagen 79216) with 1% β-mercaptoethanol (Thermo Fisher Scientific 21985023) and 5 units/ml proteinase K (New England Biolabs P8107S) at 55°C for 30 minutes. RNA and DNA were then extracted from the lysate using Trizol LS (Thermo Fisher Scientific 10296010) following the manufacturer’s instructions. RNA was subsequently converted to cDNA using iScript cDNA synthesis kit (Bio-Rad 1708890) following the manufacturer’s instructions.

### PCR and sequence analysis of tumors

*Kras* exon 1 to 3 was amplified from the cDNA of tumor or normal lung tissue using nested PCR. Primers are listed in Supplementary file 3. PCR reactions were as follows:

<u>PCR1:</u> 1 μl cDNA, 0.5 μl of 10 μM forward primer and 0.5 μl reverse primer, 2 μl of 2.5 mM dNTP (New England Biolabs N0447S), 0.75 μl of 50 mM MgCl_2_, 2 μl of 5X buffer, and 0.1 μl Platinum Taq Polymerase (Thermo Fisher Scientific 10966083) in a total volume of 25 μl. PCR cycles were as follows: one cycle at 94°C for 2 min and 25 cycles at 94°C for 30 seconds, 56°C for 30 seconds, 72°C for 30 seconds.
<u>PCR2:</u> 2.5 μl of PCR1 reaction, 0.5 μl of 10 μM forward primer and 0.5 μl reverse primer, 2 μl of 2.5 mM dNTP, 0.75 μl of 50 mM MgCl_2_, 2 μl of 5X buffer, and 0.1 μl Platinum Taq Polymerase in a total volume of 25 μl. PCR cycles were as follows: one cycle at 94°C for 2 min and 20 cycles at 94°C for 30 seconds, 58°C for 30 seconds, 72°C for 35 seconds.

10 μl product for PCR2 was then analyzed with gel electrophoresis to check PCR efficiency. For samples with no product, *Kras* exon 1 and 2 was amplified from the DNA of the same tumor separately using nested PCR. Primers were listed in Supplementary file 3. PCR reactions were as follows:

<u>PCR1:</u> 1 μl cDNA, 0.5 μl of 10 μM forward primer and 0.5 μl reverse primer, 2 μl of 2.5 mM dNTP, 0.75 μl of 50 mM MgCl_2_, 2 μl of 5X buffer, and 0.1 μl Platinum Taq Polymerase in a total volume of 25 μl. PCR cycles were as follows: one cycle at 94°C for 2 min and 25 cycles at 94°C for 30 seconds, 53°C for 30 seconds, 72°C for 15 seconds.
<u>PCR2:</u> The reactions were comprised of 5 μl PCR1, 0.5 μl of 10 μM forward primer and 0.5 μl reverse primer, 2 μl of 2.5 mM dNTP, 0.75 μl of 50 mM MgCl_2_, 2 μl of 5X buffer, and 0.1 μl Platinum Taq Polymerase in a total volume of 25 μl. PCR cycles were as follows: one cycle at 94°C for 2 min and 20 cycles at 94°C for 30 seconds, 57°C for 30 seconds, 72°C for 18 seconds.

Products from PCR2 were pooled and purified with Ampure XP beads according to the manufacturer’s protocol (Beckman Coulter A63880). The library was sequenced using MiSeq v2 Nano 250 PE at Duke Center for Genomic and Computational Biology. All primers were synthesized by Integrated DNA Technologies.

### Sequencing data analysis

Raw sequencing data were uploaded to usegalaxy.org (Jalili et al., 2020). For analysis of amplicon from cDNA and exon 2 from genomic DNA, paired-end reads were joined with fastq-join tool, with the maximum percentage difference between matching segments set at 20% and the minimum length of matching segments set at 10. For analysis of amplicon from exon 1 from genomic DNA, only read 1 was used. The joined-read for cDNA and exon 2, or read 1 for exon 1, were then processed with Filter by Quality Tool, requiring 75% of bases have quality equal to or higher than the cut-off value of 20. For the analysis of amplicon from cDNA, the forward and reverse index was then extracted from the filtered reads through the Trim Tool. Bases encoding codons 12, 13, 61, as well as nucleotide 96 (where a SNP exists that differentiates 129 from B6 strains) and 8 nucleotides in the middle of exon 3 (where the sequence differs between native and ex3op versions of *Kras*) were appropriate were also extracted through the Trim Tool. Collapsing by the index (representing individual samples) and counting the reads for each variant of the extracted region-of-interest were then performed by the Group Tool. Mutation and allele information were then assigned to each extracted region-of-interest by comparing the sequences of the extracted region-of-interest with reference sequences for all possible missense mutations in codon 12, 13, 61, SNP at nucleotide 96, and either native or codon-optimized exon 3 using the Excel program. The fraction of each variant of the extracted region was calculated by dividing the counts for that variant by the total number of counts per sample. For analysis of amplicon from genomic DNA, quality filtered-reads were split into separate files using the Barcode Splitter Tool and 5’ index. Each of the files were then trim from 3’ end to expose the 3’ index. Trimmed files were further split into separate files corresponding to different samples using the Barcode Splitter Tool and 3’ index. For each sample, bases covering codons 12 and 13 in exon 1 and codon 61 in exon 2 were extracted through the Trim Tool. The mutation was assigned to extracted bases by comparing them against reference sequences for all possible missense mutations in codon 12, 13, and 61 using the Excel program. The fraction of each variant was calculated by dividing the counts for that variant by the total number of counts per sample. Samples with less than 30 total reads or variants with fraction less than 8% were excluded from analysis.

### qPCR analysis of *Trp53* recombination

Quantitative PCR (qPCR) reactions were performed using iTaq Universal SYBR Green Supermix (Bio-Rad 1725120) and CFX384 touch real-time PCR detection system (Bio-Rad). Reaction were comprised of 200 ng gDNA, 5 μl Supermix, 0.5 μl forward primer, 0.5 μl reverse primer in a total volume of 10 μl. PCR conditions were one cycle at 95°C for 3 minutes, 40 cycles at 95°C for 10 seconds, 55°C for 30 seconds, and a melt curve cycle (65-95°C at 0.5°C increments at 5 seconds/step). Primers were designed to detect unrecombined *Trp53^flox^* allele (p53 WT), recombined *Trp53^flox^* allele (p53 Del) and the reference gene *Tflc*. Primer sequences were:

*Trp53* WT and Del F: 5’-ATCCTTTATTCTGTTCGATAAGCTTG-3’
*Trp53* WT R: 5’-AGGACTACACAGAGAAACCCT-3’
*Trp53* Del R: 5’-GCTATTGTAGCTAGAACTAGTGGAT-3’
*Tflc* F: 5’-ACCAAATGGTTCGTACAGCA-3’
*Tflc* R: 5’-ATGACAGTAGTTTGCTGTTATACATC-3’

The relative levels of *Trp53* WT and *Trp53* Del genomic DNA were calculated using ΔCt method in comparison to *Tflc*. The fraction of *Trp53* Del was then calculated by dividing the relative level of *Trp53* Del by the sum of the relative level of *Trp53* WT and *Trp53* Del.

### qPCR analysis of *Kras* copy number

qPCR reactions were performed using PrimeTime Gene Expression Master Mix (Integrated DNA Technologies 1055770) and CFX384 touch real-time PCR detection system (Bio-Rad). The reaction were comprised of 200 ng gDNA, 5 μl master mix, and 0.5 μl forward primer, 0.5 μl reverse primer, 0.25 μl probe for *Kras^nat^* allele, *Kras^ex3op^* allele, and reference gene *Tert* in a total volume of 10 μl. The conditions were one cycle at 95°C for 3 minutes, 40 cycles at 95°C for 10 seconds, 57°C for 30 seconds, and a melt curve (65-95°C at 0.5°C increments at 5 sec/step). Primers sequences were:

*Kras^nat^* F: 5’-GGAATAAGTGTGATTTGCCTTCT-3’
*Kras^nat^* R: 5’-ACCTGTCTTGTCTTTGCTGA-3’
*Krasex3op* F: 5’-AAGTGCGACCTCCCTAGC-3’
*Krasex3op* R: 5’-CTGTCTTGTCTTGGCGCT-3’
*Tert* F: 5’-CCTGACCATCTGGTGACAC-3’;
*Tert* R: 5’-GTGCCTTCTCAGAGAACACA-3’.

Probe sequences were:

*Kras^nat^* 5’-/5Cy5/AACAGTAGA/TAO/CACGAAACAGGCTCAGGA/3IAbRQSp/-3’
*Krasex3op:* 5’-/5HEX/AACCGTGGA/ZEN/CACCAAGCAGGCC/3IABkFQ/-3’
*Tert:* 5’-/56-FAM/TGGAACCAA/ZEN/ACATACATGCAGGTGCAG/3IABkFQ/-3’.

The relative levels of *Kras^nat^* and *Kras^ex3op^* alleles were calculated using ΔCt method in comparison to *Tert*. The copy number of *Kras^nat^* and *Kras^ex3op^* allele were then determined by comparing the relative DNA level in tumor samples to normal lung tissues from *Kras^nat/nat^* or *Kras^ex3o/ex3op^* mice, which were used as reference for two copies of *Kras^nat^* or *Kras^ex3op^* allele, respectively.

### qPCR analysis of MAPK signaling

qPCR reactions were performed using iTaq Universal SYBR Green Supermix (Bio-Rad 1725120) and CFX384 touch real-time PCR detection system (Bio-Rad). The reaction were comprised of 1 μl gDNA, 5 μl Supermix, 0.5 μl forward primer, 0.5 μl reverse primer in a total volume of 10 μl. The conditions were one cycle at 95°C for 3 minutes, 40 cycles at 95°C for 10 seconds, 58°C for 30 seconds, and a melt curve (65-95°C at 0.5°C increments at 5 sec/step). The primers sequences were:

*Dusp6* F: 5’-ACTTGGACGTGTTGGAAGAGT-3’
*Dusp6* R: 5’-GCCTCGGGCTTCATCTATGAA-3’
*Egr1* F: 5’-CCTGACCACAGAGTCCTTTTCT-3’
*Egr1* R: 5’-AGGCCACTGACTAGGCTGA-3’
*Fosl1* F: 5’-CAGGAGTCATACGAGCCCTAG-3’
*Fosl1* R: 5’-GCCTGCAGGAAGTCTGTCAG-3’
*Actin* F: 5’-CGTGAAAAGATGACCCAGATCATGT-3’
*Actin* R: 5’-CGTGAGGGAGAGCATAGCC-3’.

Gene expression values were calculated using the comparative Ct (-ΔΔCt) method (Livak and Schmittgen, 2001), using actin as internal control.

### Isolation of genomic DNA for maximum depth sequencing assay

Lung tissues were cut into fine pieces and resuspended in 500 μl lysis buffer (100 mM NaCl, 10 mM Tris pH 7.6, 25 mM EDTA pH 8.0, and 0.5% SDS in H_2_O, supplemented with 20 μg.ml^−1^ RNase A (Sigma R4642). Samples were incubated at 37°C for 1 hour. 2.5 μl of 800 U.ml^−1^ proteinase K (New England Biolabs P8107S) was then added to each sample, then the samples were vortexed and incubated at 55°C overnight. Genomic DNA was isolated by phenol/chloroform extraction followed by ethanol precipitation using standard procedures.

### Maximum depth sequencing (MDS)

This method was adapted from published protocols (Jee et al., 2016; Li et al., 2020). In detail, 20-50 μg of genomic DNA was incubated with EcoRV (New England Biolabs R3195), EcoRI (New England Biolabs R3101), and XmnI (New England Biolabs R0194) for the analysis of the non-transcribed strand of *Kras* exon 1 and exon 2. Reaction conditions were 5 units of the each of the indicated restriction enzymes and per 1μg DNA per 20 μl reaction. Digested genomic DNA was column purified using QIAquick PCR Purification Kit following the manufacturer’s protocol (Qiagen 28104) and resuspended in ddH_2_O (35 μl H_2_O per 10 μg DNA). The barcode and adaptor were added to the target DNA by incubating purified DNA with the appropriate barcode primers (*see* below) for one cycle of PCR. PCR reactions were comprised of 10 μg DNA, 2.5 μl of 10 μM barcode primer (*see* below), 4 μl of 2.5 mM dNTP, 10 μl of 5X buffer, and 0.5 μl Q5^®^ Hot Start High-Fidelity DNA Polymerase (New England Biolabs M0493) in a total volume of 50 μl. The number of PCR reactions was scaled according to the amount of DNA. PCR conditions were 98°C for 1 minute, 60°C for 15 seconds, and 72°C for 1 minute using the barcoding primer for exon 2, followed by the addition of the barcoding primer for exon 1 to the same reaction, 98°C for 1 minute, 68°C for 15 seconds, and 72°C for 1 minute. 1 μl of 20,000 U.ml^−1^ exonuclease I (New England Biolabs M0293) and 5 μl of 10X exonuclease I buffer (New England Biolabs B0293S) was then added to each 50 μl reaction to remove unused barcoded primers and incubated at 37°C for 1 hour and then 80°C for 20 minutes. Processed DNA were column-purified using QIAquick PCR Purification Kit as above and resuspended in ddH_2_O (35 μl H_2_O per column). The concentration of purified product was measured with SimpliNano spectrophotometer (GE Healthcare Life Sciences). Samples were linear amplified with forward adaptor primer (*see* below). PCR reactions were comprised of 1.5 μg DNA, 2.5 μl of 10 μM forward-adaptor primer, 4 μl of 2.5 mM dNTP, 10 μl of 5X buffer, and 0.5 μl Q5^®^ Hot Start High-Fidelity DNA Polymerase in a total volume of 50 μl. The number of PCR reactions was scaled according to the amount of DNA. PCR conditions were as follows: 12 cycles of 98°C for 15 seconds, 70°C for 15 seconds, and 72°C for 10 seconds. 2.5 μl of 10 μM exon-specific reverse primers (*see* below) and 2.5 μl of 10 μM reverse-adaptor primer (*see* below) were then added to each 50 μl reaction. The mixtures were then subjected exponential amplification. PCR conditions were as follows: 4 cycles of 98°C for 15 seconds, 62°C for 15 seconds, 72°C for 10 seconds, 20 cycles of 98°C for 15 seconds, 70°C for 15 seconds, and 72°C for 10 seconds. The final library was size selected and purified with Ampure XP beads according to the manufacturer’s protocol (Beckman Coulter A63880). Sequencing was performed using NovaSeq 6000 S Prime 150bp PE at Duke Center for Genomic and Computational Biology.

### Primers for MDS

Barcode primer: [Forward adaptor][Index][Barcode][Primer]

Where

[Forward adpator] =

5’-TACGGCGACCACCGAGATCTACACTCTTTCCCTACACGACGCTCTTCCGATCT-3’

[Index] = variable length of known sequences from 0 to 7 nucleotides (Supplementary file 3)

[Barcode] = NNNNNNNNNNNNNN

*Kras* exon 1 [Primer] = 5’-ATCTTTTTCAAAGCGGCTGGCT-3’

*Kras* exon 2 [Primer] = 5’-TCTTCAAATGATTTAGTATTATTTATGGC-3’

Forward-adaptor primer: 5’-AATGATACGGCGACCACCGAGAT-3’

Exon-specific reverse primer: [Reverse adaptor][Index][Primer]

Where

[Reverse adaptor] =

5’-CAAGCAGAAGACGGCATACGAGATGTGACTGGAGTTCAGACGTGTGCTCTTCCTCT-3’

[Index] = variable length of known sequences from 0 to 7 nucleotides (Supplementary file 3)

*Kras* exon 1 [Primer] = 5’-TATTATTTTTATTGTAAGGCCTGCTGA-3’

*Kras* exon 2 [Primer] = 5’-GACTCCTACAGGAAACAAGT-3’

Reverse-adaptor primer: 5’-CAAGCAGAAGACGGCATACGAGA-3’

All primers were synthesized by Integrated DNA Technologies.

### Analysis of MDS data

Raw data were uploaded Galaxy Cloudman (Jalili et al., 2020). Read 1 and read 2 were joined via PEAR pair-end read merger (Zhang et al., 2014b). The reads were then filtered by quality by requiring 90% of bases in the sequence to have a quality core ≥ 20. Filtered reads were split into different files based on assigned sample indexes and variation in sequence lengths using the Barcode Splitter tool and Filter sequences by length tool. The reads were trimmed down to the barcode and the target exon. Trimmed reads were grouped by barcode. Barcode families containing ≥ 2 reads and have ≥ 90% reads being identical are selected. Sequences from selected barcode families were compared against annotated reference mutant sequences containing all possible single nucleotide substitutions in the exon of interest and the mutation in the reference mutant sequence was assigned to the matched barcode family. The frequency of the corresponding mutation was calculated by dividing the counts of the families containing the mutation by the total number of families.

### Generation of heatmaps

All heatmaps were generated using Morpheus (https://software.broadinstitute.org/morpheus). The mutation frequencies used in heatmaps in ***Figure 3*** and ***Figure 3-figure supplement 1*** were corrected by the addition of 1 x 10^-5^ (the detection limit at a barcode recovery of 1 x 10^5^), log_10_ transformed and plotted.

### Statistics

The number of independent experiments and the statistical analysis used are indicated in the legends of each figure. Data are represented as mean ± SEM. *p* values were determined by Dunn’s multiple comparison test following Kruskal-Wallis test, Holm-Sidak multiple comparisons test following one-way ANOVA, two-tailed Mann-Whitney U test, or two-sided Fisher’s exact test. For correlation analysis, *Rho* and *p* values were derived from the Spearman correlation test. All statistical test were performed using GraphPad Prism 6.

## Data Availability

All raw sequencing data has been deposited to NCBI Sequence Read Archive (SRA) under accession number PRJNA663179.

## Acknowledgements

We thank the head of and/or members of the laboratories of Drs. James Alvarez, Christopher Counter, David MacAlpine, and Nikoleta Tsvetanova (Duke University) for thoughtful discussions. This work was supported by the National Cancer Institute (R01CA94184 and P01CA203657 to CMC).

## Author Contributions

SL performed while CMC oversaw all experiments, both authors contributed to the writing of the manuscript.

---

**Figure 1-figure supplement 1.**
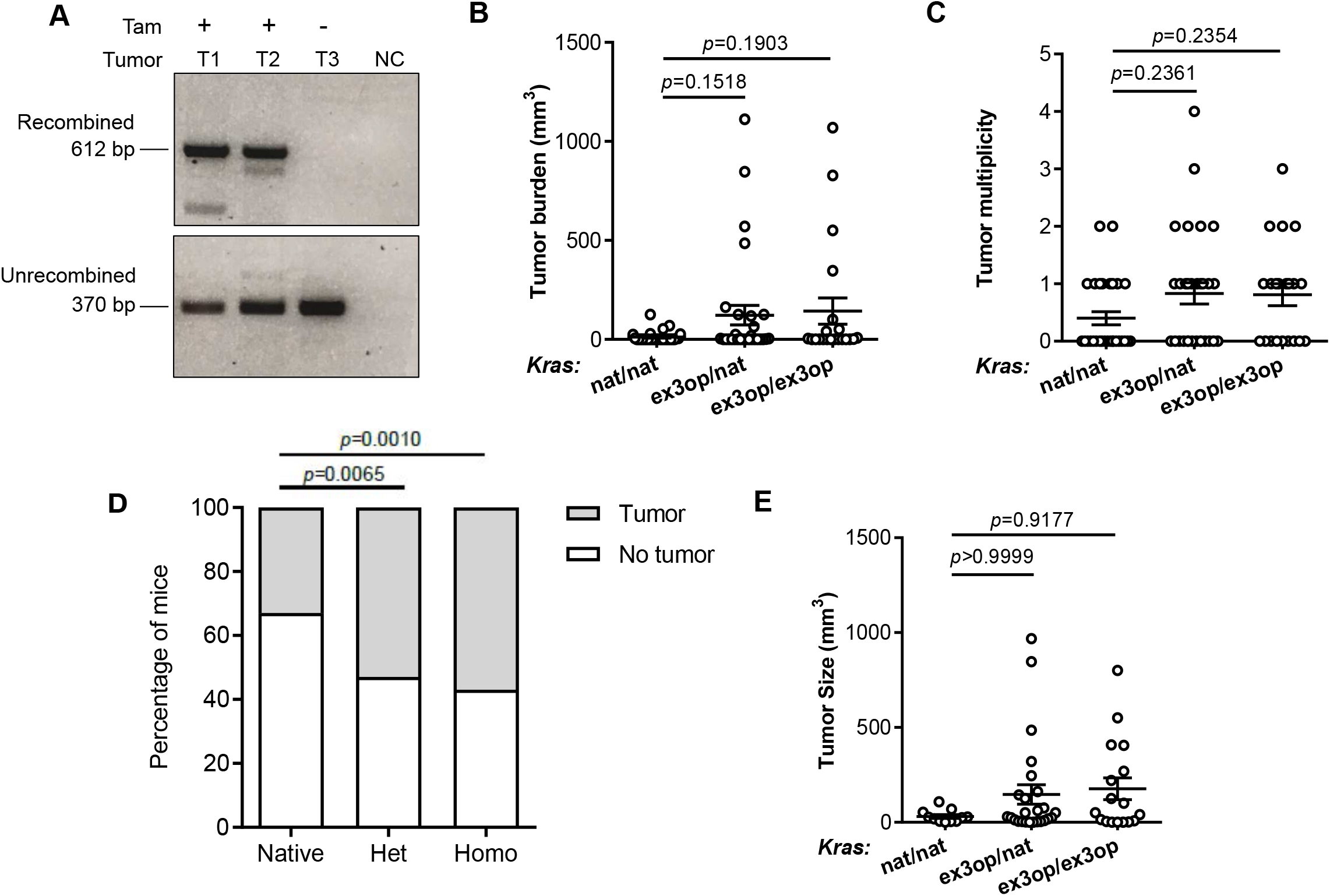
The effect of *Kras^ex3op^* allele on urethane-mediated lung tumorigenesis in the absence of p53. (**A**) PCR analysis of the status of the *Trp53^flox^* allele in DNA isolated from lung tumors in *Sftpc^CreER/CreER^;Trp53^fl/fl^;Kras^ex3op/nat^* (T1,T3) or *Sftpc^CreER/CreER^;Trp53^fl/fl^;Kras^ex3op/ex3op^* (T2) mice treated (+) or not treated (-) with tamoxifen. NC: no DNA control. (**B**, **C**, and **E**) Mean ± SEM of urethane-induced tumor (**B**) burden, (**C**) multiplicity, and (**E**) size in tamoxifen-treated *Sftpc^CreER/ CreER^;Trp53^fl/fl^* mice in a homozygous native ((**B** and **C**) n=30 mice, (**E**) n=11 tumors), heterozygous ((**B** and **C**) n=30 mice, (**E**) n=25 tumors) and homozygous ((**B** and **C**) n=21 mice, (**E**) n=17 tumors) ex3op Kras background. Dunn’s multiple comparison test following Kruskal-Wallis test. (**D**) % of mice with (grey bar) or without (white bar) a tumor in tamoxifen-treated *Sftpc^CreER/CreER^;Trp53^fl/fl^* mice in *Kras^nat/nat^* (n=30 mice), *Kras^ex3op/nat^* (n=30 mice), or *Kras^ex3op/ex3op^* (n=21 mice) background after urethane exposure. Two-sided Fisher’s exact test.

**Figure 2-figure supplement 1.**
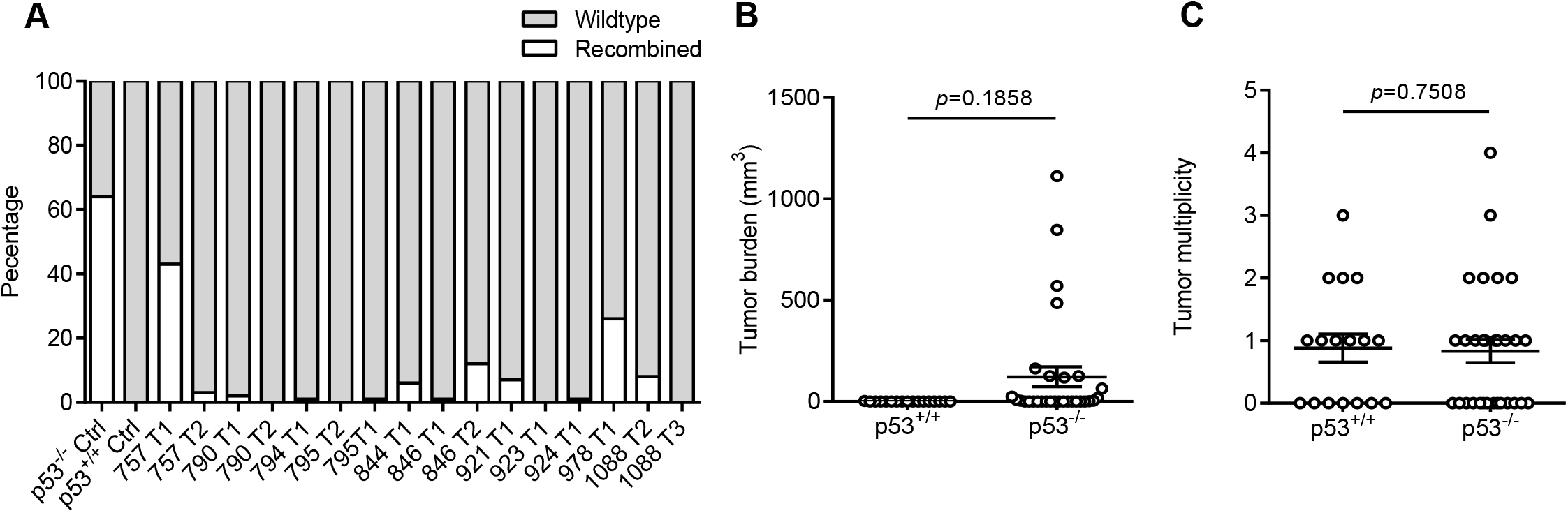
The effect of p53 loss on tumor burden and multiplicity. (**A**) Proportion of recombined p53 allele and widltype allele in tumors from mice not treated with tamoxifen by qPCR. (**B**,**C**) Mean ± SEM tumor (**B**) burden and (**C**) multiplicity of *Sftpc^CreER/CreER^;Trp53^fl/fl^;Kras^ex3op/nat^* mice not treated (p53^+/+^, n=17 mice) or treated (p53^-/-^, n=30 mice) with tamoxifen. Mann-Whitney test.

**Figure 3-figure supplement 1.**
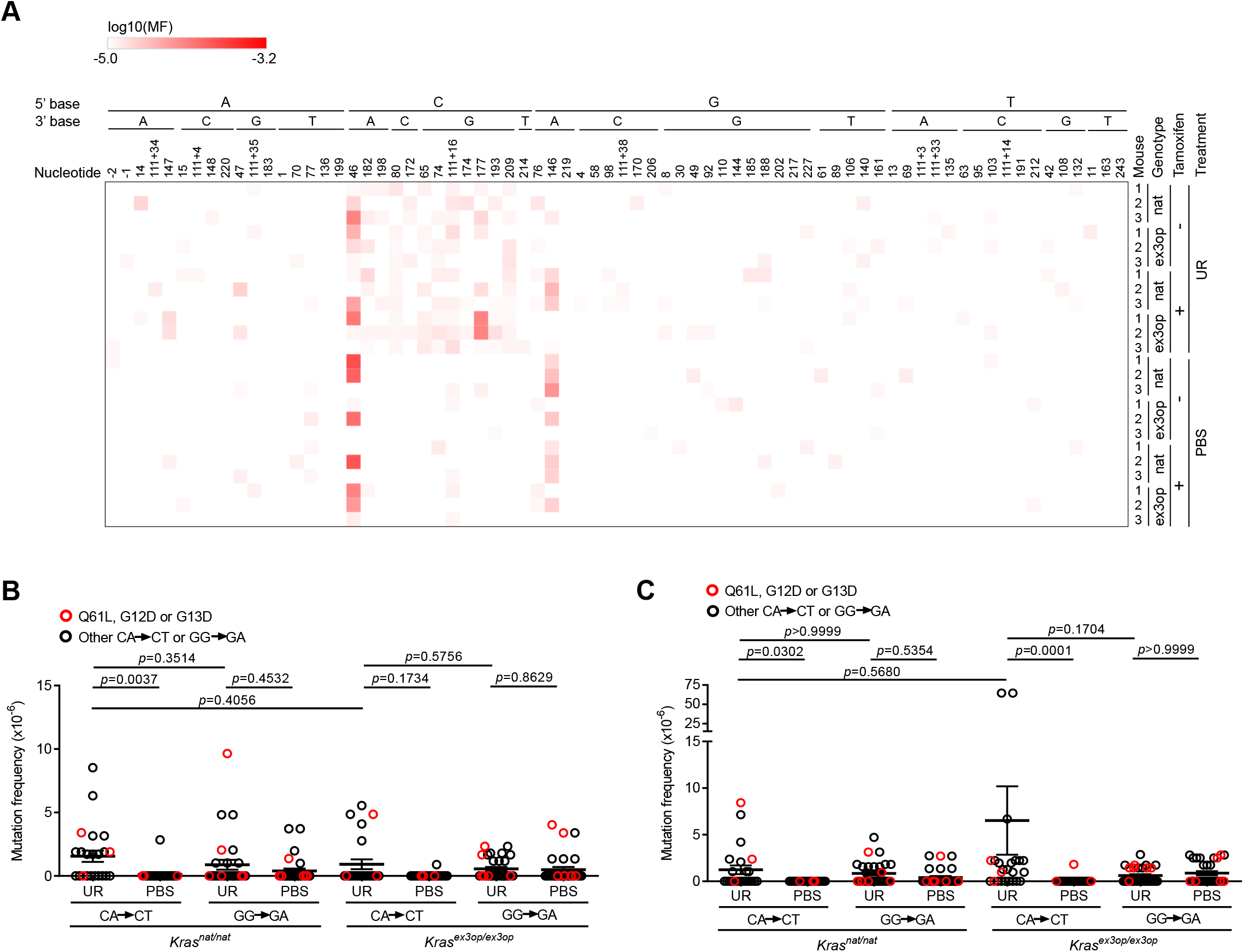
Mutagenesis profile of *Sftpc^CreER/CreER^;Trp53^flox/flox^;Kras^nat/nat^* and *Sftpc^CreEE/CreER^;Trp53^flox/flox^;Kras^ex3op/ex3op^* mice. (A) Heatmap of the log transformed mutation frequency (MF) determined by MDS sequencing the exon 1 and exon 2 of *Kras* in from the lungs of mice treated (+) or not treated with tamoxifen (-), exposed to urethane (UR) or PBS, in a *Sftpc^CreER/CreER^;Trp53^fl/ fl^;Kras^nat/nat^* (nat) or *Sftpc^CreER/CreER^;Trp53^fl/fl^;Kras^ex3op/ex3op^* (ex3op) background (n=3 mice) for each A>T transversions (nucleotide number as well as the 5’ and 3’ base of the substituted A are shown at the top, indicates nucleotides upstream of ATG start codon in 5’UTR, “111+” indicates nucleotides in the intron downstream of exon 1). **(B,C)** Mean ± SEM mutation frequency of all CA>CT mutations in *Kras* exon 2, including Q61L mutation highlighted in red, as well as all GG>GA mutations in *Kras* exon 1, including G12D and G13D mutations highlighted in red, derived from MDS sequencing of *Kras* exon 1 and 2 from the lungs of *Sftpc^CreER/CreER^;Trp53^fl/fl^;Kras^nat/nat^* versus *Sftpc^CreER/CreER^;Trp53^fl/fl^;Kras^ex3op/ex3op^* mice (B) not treated or (C) treated with tamoxifen and exposed to either urethane or PBS (n=3 mice). (B) Holm-Sidak multiple comparisons test following one-way ANOVA. (C) Dunn’s multiple comparison test following Kruskal-Wallis test.

**Figure 4-figure supplement 1.**
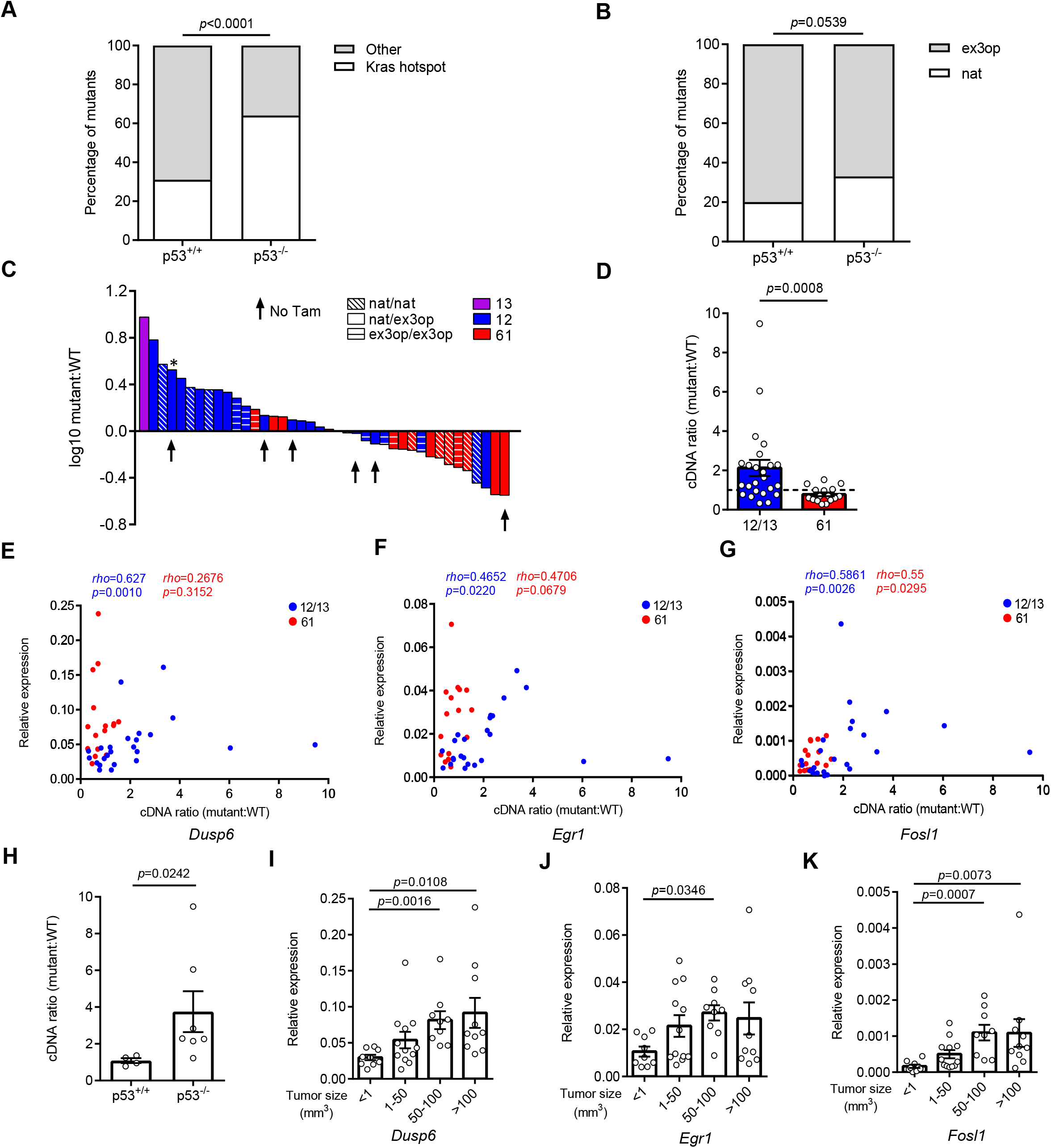
Allelic imbalance and MAPK signaling in Kras hotspot-mutant tumors. (**A**) % of tumors with an oncogenic mutation at Kras hotspot (white bar) versus other tumors (grey bar) from *Sftpc^CreER/CreER^;Trp53^fl/ fl^;Kras^ex3op/nat^* mice not treated (p53+/+, n=16 tumors) or treated (p53-/-, n=25 tumors) with tamoxifen. Two-sided Fisher’s exact test. (**B**) % of tumors with Kras hotspot mutations occurring in the native (white bar) or ex3op (grey bar) allele from *Sftpc^CreER/CreER^; Trp53^fl/fl^;Kras^ex3op/nat^* mice not treated (p53^+/+^, n=5 tumors) or treated (p53^-/-^, n=15 tumors) with tamoxifen. Two-sided Fisher’s exact test. (**C**) Log10-transformed ratio of mutant to wildtype *Kras* mRNA determined by RT-qPCR in all Kras hotspot-mutant tumors (n=40 tumors) derived from Figures 1 and 2. Asterisk indicates tumor 757T1 with a p53 deficiency. (**D**) Mean ± SEM ratio of mutant to wildtype *Kras* mRNA in tumors with G12/13 (n=24 tumors) and Q61 (n=16 tumors) mutations. (**E-G**) Correlation between the levels of the indicated mRNAs normalized to ß-actin (Relative expression) and the ratio of mutant to wildtype *Kras* mRNA. Rho and p values are from Spearman correlation analysis. Mann-Whitney test. (H) Mean ± SEM ratio of G12/13 mutant *Kras^ex3op^* to wildtype *Kras^nat^* mRNA in tumors from *Sftpc^CreER/CreER^;Trp53^fl/fl^;Kras^ex3op/nat^* mice not treated (p53^+/+^, n=4 tumors) or treated with tamoxifen (p53^-/-^, n=7 tumors). Mann-Whitney test. **(I-K**) Mean ± SEM levels of the indicated mRNAs normalized to ß-actin (Relative expression) for tumors that are <1 (n=9 tumors), 1-50 (n=12 tumors), 50-100 (n=9 tumors), and >100 (n=10 tumors) mm^3^. Dunn’s multiple comparison test following Kruskal-Wallis test.

**Figure 4-figure supplement 2.**
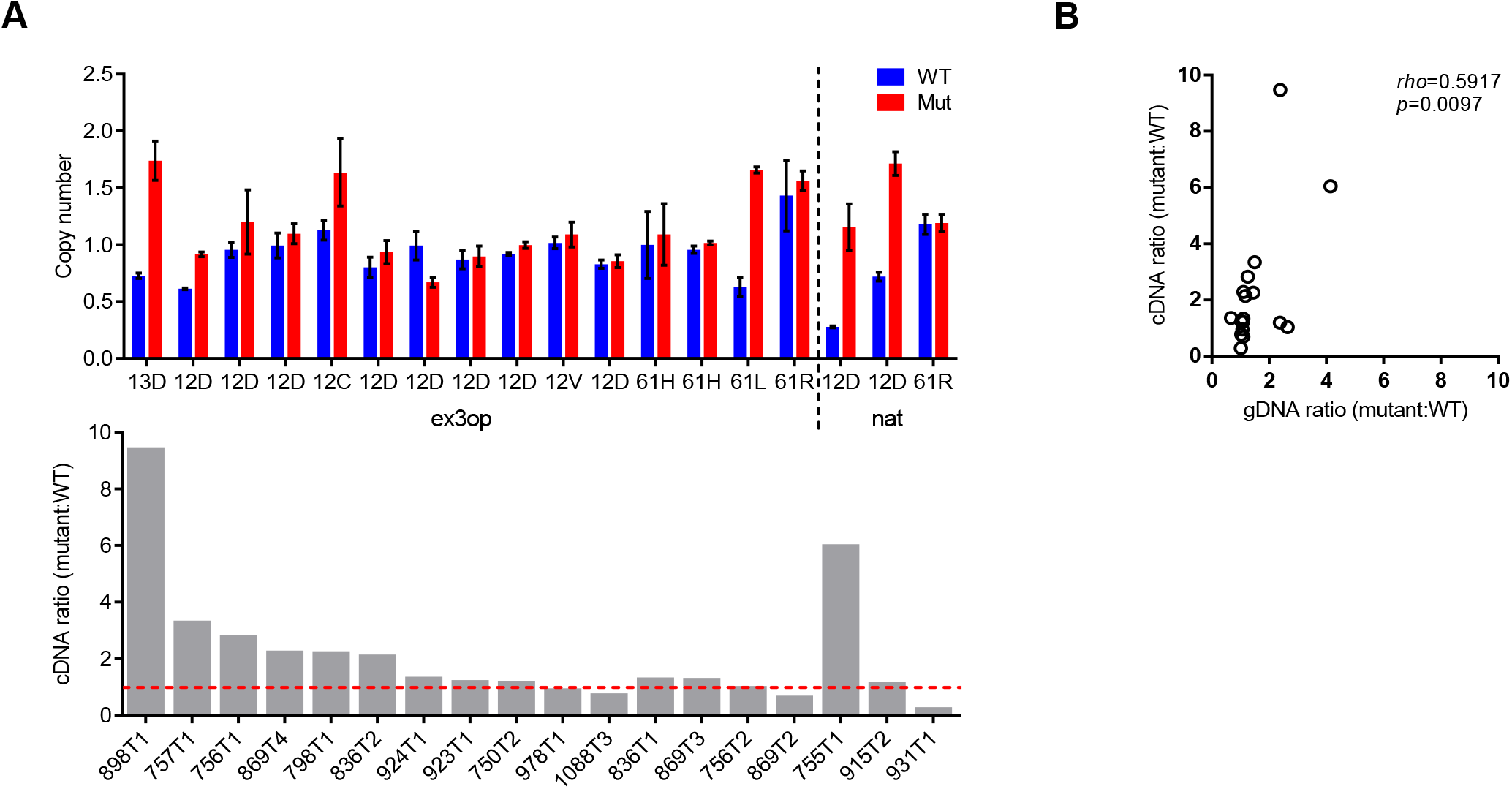
The imbalance at mRNA level could not be fully attributed to the imbalance of DNA copy number. (**A**) Top: Copy number of the mutant or wildtype allele in tumors from *Sftpc^CreER/CreER^;Trp53^fl/fl^;Kras^ex3op/nat^* mice estimated by qPCR copy number assay (Tert as reference gene). Mutant type and the allele with mutation are indicated. Bottom: the ratio of mutant to wildtype *Kras* mRNA and ID of the tumors listed in the top graph. Data shown are mean ± SEM of two technical replicates. (**B**) Correlation between the mRNA ratio and genomic DNA ratio of mutant to wildtype *Kras* allele. Rho and p values are from Spearman correlation analysis.

